# Calmodulin controls the tempo of HSV-1 gene-expression cascade to reshape infection heterogeneity

**DOI:** 10.64898/2026.06.02.729706

**Authors:** Yuhei Maruzuru, Yuta Kuze, Masahide Seki, Akinori Kanai, Shaocong Liu, Akihisa Kato, Yutaka Suzuki, Yasushi Kawaguchi

**Author notes:** Address correspondence to: Dr. Yasushi Kawaguchi, Division of Molecular Virology, Department of Microbiology and Immunology, The Institute of Medical Science, The University of Tokyo, 4-6-1 Shirokanedai, Minato-ku, Tokyo 108-8639, Japan.

## Abstract

Viral replication exhibits substantial heterogeneity among individual cells. Although temporally ordered gene-expression cascades are thought to contribute to such heterogeneity in DNA viruses, whether and how their tempo is regulated, and how cell-to-cell variability shapes population-level infection outcomes, remain unknown. Here, we developed a high-throughput live-cell imaging platform to track HSV-1 gene-expression dynamics in individual cells and, in combination with single-cell RNA sequencing, identified calmodulin (CaM) as a host regulator of HSV-1 gene-expression tempo. CaM accelerates progression from immediate-early to early and late phases at the single-cell level by advancing their onset. By increasing the fraction of virus-producing cells without altering burst size, CaM reshapes the distribution of infection outcomes, thereby promoting population-level viral replication. Pharmacological inhibition of CaM protects mice from lethal infection, exceeding acyclovir. These findings establish CaM-mediated temporal control as a key determinant of population-level viral output and a regulator of infection heterogeneity.

## INTRODUCTION

DNA viruses typically achieve their replication through a hierarchical cascade of gene expression, in which distinct temporal classes of genes are expressed sequentially ^1^. This hierarchical organization represents a fundamental strategy to maximize replication efficiency by ensuring that enzymes and regulatory factors required for DNA synthesis are produced prior to the resource-intensive synthesis of structural components, thereby optimizing viral propagation within the host cell ^1^. At the same time, viral replication is known to exhibit substantial heterogeneity among individual infected cells, and this hierarchical organization of viral gene expression likely contributes to diverse infection outcomes ^2–6^. However, the molecular mechanisms that govern the tempo and fidelity of progression through this hierarchical cascade within individual cells—and even whether this tempo is actively regulated—have remained largely unexplored.

Despite the central importance of temporal regulation, two fundamental questions remain unresolved: (i) whether specific host determinants exist that tune the tempo of the viral gene expression cascade, and (ii) how single-cell temporal dynamics shape viral production at the population level. Addressing these questions has been hindered by technical limitations. Bulk assays mask cell-to-cell heterogeneity ^7,8^, whereas single-cell RNA sequencing provides only static snapshots and lacks temporal resolution ^9^. Although time-lapse imaging can capture viral gene expression dynamics in individual cells ^10,11^, robust single-cell segmentation and long-term tracking in confluent cultures remain significant challenges. Consequently, most analyses are restricted to low cell density condition that fail to recapitulate the cell-cell contacts characteristic of physiological environments. Crucially, no quantitative framework currently links phenotypic diversity at the single-cell level to total viral yield, leaving the causal relationship between single-cell heterogeneity and population-level replication unresolved.

To address these questions, we first sought to identify host factors associated with progression through the viral gene-expression cascade. We focused on herpes simplex virus type 1 (HSV-1), a prototypic large DNA virus of the *Herpesviridae* family, in which viral genes are expressed in a temporally ordered cascade comprising immediate-early (IE), early (E), and late (L) classes ^12^. Using single-cell RNA sequencing (scRNA-seq) of HSV-1–infected cells, we identified *CALM1* and *CALM2* as host genes whose expression levels positively correlated with viral transcript abundance. These genes encode calmodulin (CaM), a ubiquitous calcium sensor known to regulate critical cellular functions, including transcriptional processes ^13–16^.

Motivated by this finding, we developed a fully automated, high-throughput imaging workflow that enables long-term tracking of viral gene expression in individual cells at standard culture densities. This system achieves quantitative accuracy comparable to flow cytometry while allowing direct visualization of temporal kinetics at the single-cell level. By integrating this imaging workflow with cell sorting–based analyses that quantitatively link single-cell gene expression to population-level virus production, we demonstrate that CaM, through its interaction with the major HSV-1 transcriptional regulator ICP4 ^12,17^, promotes progression from IE to E and L phases at the single-cell level by advancing their onset, thereby regulating the tempo of the hierarchical cascade. This regulation reshapes the distribution of infection outcomes by increasing the fraction of virus-producing cells without altering their burst size, leading to promotion of progeny viral production at the population level.

Our findings identify a previously unrecognized regulatory layer in the canonical viral gene-expression cascade and reveal a host mechanism linking single-cell gene-expression dynamics to population-level viral replication, providing a framework for understanding how infection heterogeneity is shaped.

## RESULTS

### CALM1 and CALM2 expression are associated with the progression through HSV-1 gene-expression cascade

A previous scRNA-seq study suggested that individual HSV-1–infected cells exhibit substantial variability in viral load and that this heterogeneity reflects differences in progression along the viral gene-expression cascade ^5^. These observations raised the possibility that host factors may modulate progression through the viral gene-expression cascade in single cells, thereby shaping the observed heterogeneity. To identify such host regulators, we first examined host transcripts that correlate with this progression at the single-cell level.

HeLa cells infected with wild-type HSV-1(F) at an MOI of 5 were analyzed at 13 h post-infection by single-cell RNA sequencing. Consistent with previous observations ^5,18^, the fraction of HSV-1–derived reads varied widely across individual cells (0.02–51%; S-Fig. 1A), with increasing viral read fraction associated with decreased proportions of immediate-early (IE) and early (E) expression (IE: ρ = –0.57, p < 0.0001; E: ρ = –0.48, p < 0.0001) and increased late (L) gene expression (ρ = 0.67, p < 0.0001) (S-Fig. 1B–D). These results confirm that, even under high-MOI conditions, HSV-1 gene expression is heterogeneous across individual cells and reflects differences in progression through the viral transcriptional cascade.

Because HSV-1 infection extensively alters host gene expression ^19,20^, correlations between host and viral reads may largely reflect virus-induced transcriptional changes rather than pre-existing cellular differences that influence viral gene expression. To minimize such confounding, we focused on 1,566 host genes whose expression relative to the host transcriptome was not significantly altered between infected (n = 77) and uninfected (n = 84) cells (q > 0.05, –1 < log2FC < 1) (S-Fig. 1E and F) and examined which of these correlated with the viral read fraction. Among the genes showing a significant positive correlation, CALM1 emerged as one of the strongest (Fig. 1A and B). CALM1 encodes calmodulin (CaM), a highly conserved calcium-binding protein that mediates calcium-dependent signaling ^13^. In humans, CaM is encoded by three genes (CALM1/2/3) that produce identical proteins ^21^. We therefore examined whether the expression of CALM2 and CALM3 also correlated with the viral read fraction. CALM2 showed a similar positive correlation, whereas CALM3 did not (Fig. 1C and D), likely due to its low expression level (Fig. 1E). Thus, CALM1 and CALM2 expression positively correlates with viral read fraction, a proxy for progression through the viral gene-expression cascade at the single-cell level.

**Fig. 1.**
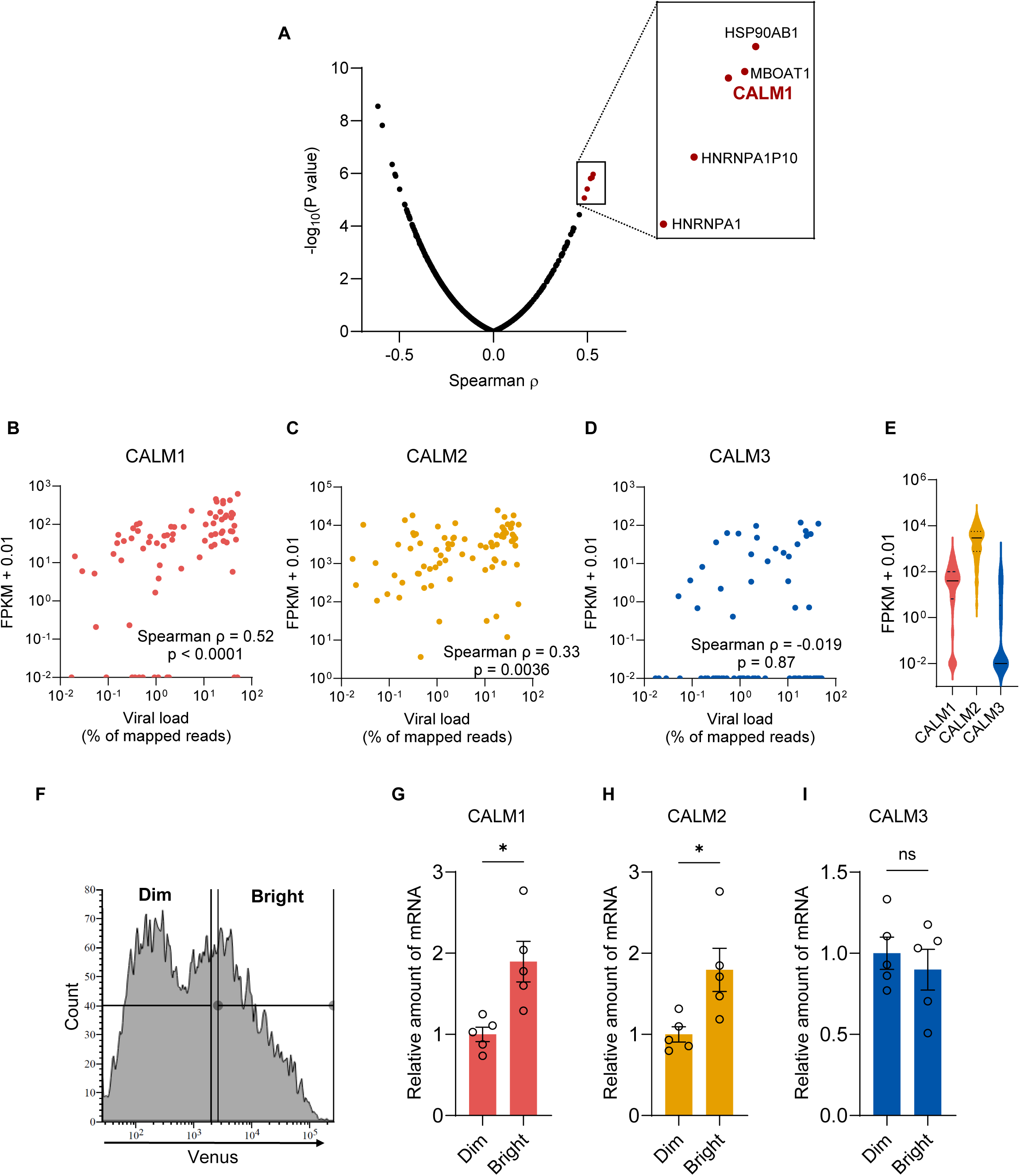
CALM1 and CALM2 expression correlates with the progression of HSV-1 gene expression. (A) Volcano plot showing Spearman’s ρ and –log₁₀(p value) for each of the 1,566 host genes whose expression relative to the host transcriptome was not significantly altered between HSV-1–infected and mock-infected cells. (B to D) Scatter plots of viral load (% of mapped reads) versus FPKM + 0.01 for CALM1 (B), CALM2 (C), and CALM3 (D) from 77 HSV-1-infected HeLa cells obtained in this study. Spearman’s rank correlation coefficients (ρ) and p values are indicated. (E) Violin plots showing the distribution of CALM1, CALM2, and CALM3 expression levels (FPKM + 0.01) among the 77 HSV-1-infected single cells. Solid lines indicate medians; dashed lines indicate quartiles. (F) Flow-cytometric sorting strategy for rICP47/vUs11 (IE: TagRFP; L: Venus)–infected HeLa cells at 12 h post-infection, separating Venus-Dim and Venus-Bright subpopulations. (G to I) Quantitative RT-PCR analysis of CALM1 (G), CALM2 (H), and CALM3 (I) mRNA levels in sorted cells. Data were normalized to GAPDH expression and are presented as mean ± SE of biological replicates (n = 5). Statistical analyses were performed using Welch’s t-test; *, p < 0.05; ns, not significant.

To validate the scRNA-seq result that higher CALM1/2 expression was associated with more progressed viral gene expression, we used rICP47/vUs11, a dual-fluorescent reporter virus in which TagRFP and Venus are fused to the IE protein ICP47 and the L protein Us11, respectively such that Venus fluorescence serves as a proxy for L gene expression ^22^. HeLa cells infected with this virus were sorted into Venus-Dim and Venus-Bright populations (Fig. 1F), and CALM1/2/3 expression was quantified by qPCR. CALM1 and CALM2 were significantly higher in Bright than in Dim cells, whereas CALM3 remained unchanged (Fig. 1G to I), confirming that cells with higher L gene expression also expressed higher levels of CALM1 and CALM2. The positive correlation between CALM1/2 expression and viral load fraction was also observed upon reanalysis of a previously published scRNA-seq dataset of HSV-1–infected human brain organoids,^23^ whereas CALM3 expression did not show a significant correlation (S-Fig.1G to I). Together, these results identify CALM1/2 as host genes whose expression is associated with progression of the HSV-1 gene-expression cascade.

### CaM deficiency impairs HSV-1 gene expression and attenuates viral replication

Given the positive correlation between CALM1/2 expression and progression through the HSV-1 gene-expression cascade revealed by scRNA-seq, we next examined whether CaM functionally contributes to viral gene expression. To this end, we generated CALM1 knockout (CALM1-KO) and CALM1/2 double knockout (CALM1/2-KO) HeLa cell lines using CRISPR-Cas9 (S-Fig. 2A and B). CaM protein levels decreased stepwise with the number of disrupted CALM genes (S-Fig. 2C and D), whereas cell viability remained comparable to that of wild-type cells (S-Fig. 2E). We then assessed HSV-1 gene expression by infecting these cells with wild-type HSV-1(F) at an MOI of 5 and quantifying transcripts of IE (ICP0, ICP27), E (ICP8, UL50), and L (UL49, Us2) genes at 6 h post-infection by absolute qPCR. IE gene expression was only modestly affected, with ICP27 showing a small but significant reduction only in CALM1/2-KO cells (1.4-fold; Fig. 2A and B), whereas other changes were not statistically significant. In contrast, E and L gene expression was significantly reduced in both CALM1-KO and CALM1/2-KO cells compared to wild-type cells, with the magnitude of reduction increasing stepwise with the number of disrupted CALM genes and exceeding that observed for IE genes (Fig. 2A and B). These results indicate that CaM preferentially promotes E and L gene expression. Of note, absolute quantification yielded results consistent with those obtained using the ΔΔCt method, with values from the two approaches showing a strong correlation (slope of approximately 1; S-Fig. 2F and G). Therefore, subsequent qPCR analyses were performed using the ΔΔCt method.

**Fig. 2.**
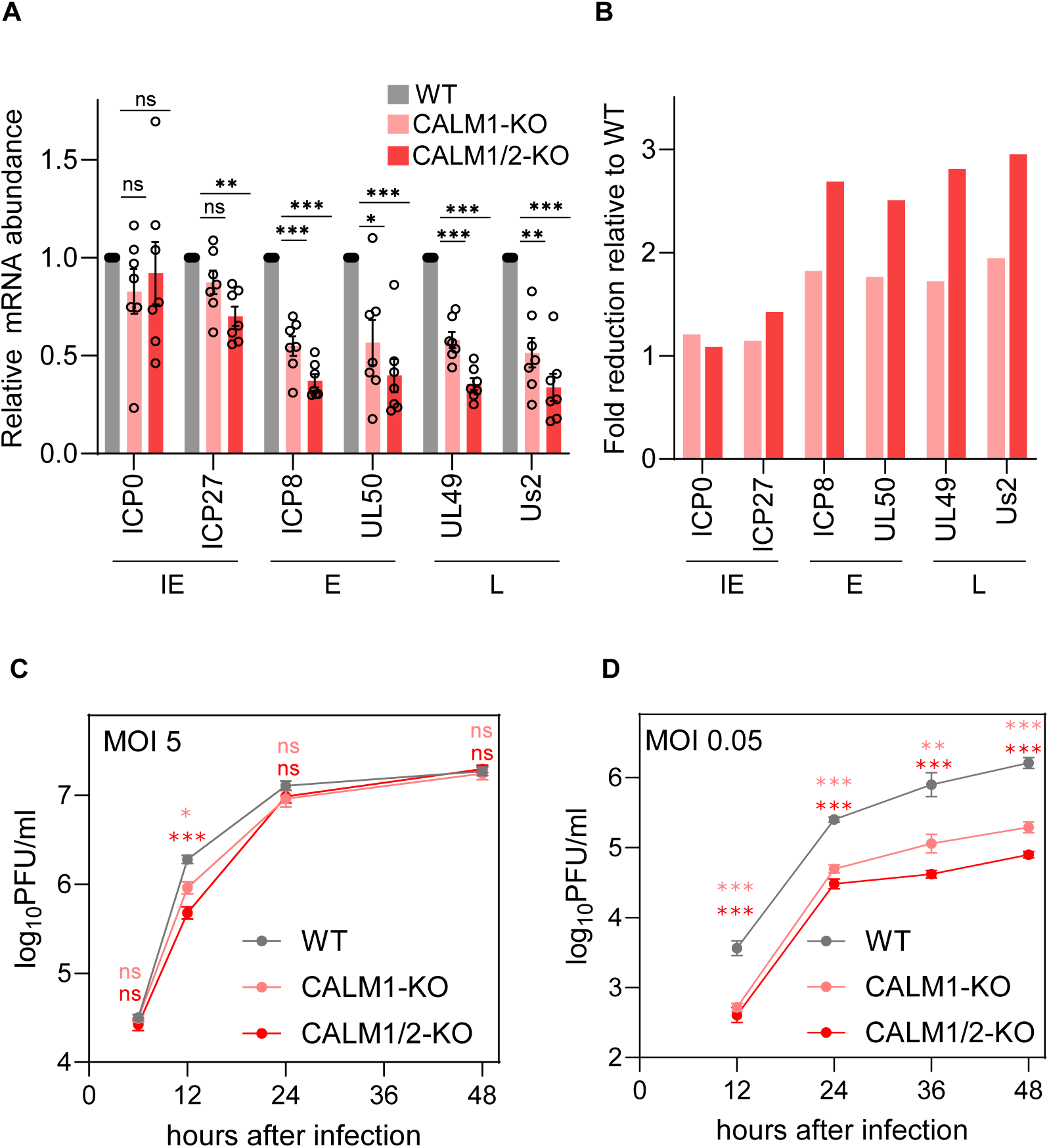
CaM deficiency impairs HSV-1 gene expression and attenuates viral replication. (A and B) Absolute quantification by RT-qPCR of representative immediate-early (IE; ICP0, ICP27), early (E; ICP8, UL50), and late (L; UL49, Us2) mRNAs in HeLa WT, CALM1-KO, and CALM1/2-KO cells at 6 h post-infection. mRNA copy numbers were normalized to the copy number of 18S rRNA. (A) mRNA abundance shown relative to WT. (B) Fold reduction relative to WT, calculated from the mean values in (A). (C and D) HeLa WT, CALM1-KO, and CALM1/2-KO cells were infected with wild-type HSV-1(F) at an MOI of 5 (C) or MOI of 0.05 (D). Each cell culture supernatant, along with the infected cells, was harvested at the indicated times post-infection, and progeny viruses were assayed on Vero cells. Each value represents the mean ± SE of seven (A) or four (C and D) biological replicates. Statistical analyses were performed using a one-sample t-test with Bonferroni correction (A) or one-way ANOVA followed by Tukey’s multiple-comparison test (C and D). In (C and D), statistical comparisons were made against WT, and the colors of the significance indicators (asterisks or ns) correspond to those of the CALM1-KO and CALM1/2-KO data. *, p < 0.05; **, p < 0.01; ***, p < 0.001; ns, not significant.

To assess the impact of CaM deficiency on HSV-1 replication, we measured progeny virus yields. Under the high-MOI condition (MOI of 5), viral titers in CaM-deficient cells were significantly reduced at 12 h post-infection but became comparable to those in wild-type cells at later time points (Fig. 2C). In contrast, under the low-MOI condition (MOI of 0.05), viral replication was significantly attenuated throughout the time course (12–48 h) (Fig. 2D). To rule out off-target effects of CRISPR-Cas9 editing, we performed rescue experiments by stably expressing Flag-tagged CaM in CALM1-KO cells (S-Fig. 2H). In CALM1-KO/Flag-CaM cells, both E/L gene expression and progeny virus yields were restored to wild-type levels (WT/puroR) (S-Fig. 2I to N), confirming that the observed phenotypes are attributable to CALM1 disruption.

Together, these results indicate that, at the population level, CaM promotes progression through the HSV-1 gene-expression cascade, particularly at the E and L stages, thereby enabling efficient HSV-1 replication.

### CaM deficiency predominantly delays the onset of E and L gene expression rather than post-onset accumulation

The preferential reduction of E and L gene expression in CaM-deficient cells at the population level suggested that progression from IE to downstream viral gene classes might be impaired in individual cells. Because this progression is inherently dynamic and heterogeneous, and cannot be resolved by static or bulk measurements, direct analysis requires time-resolved single-cell imaging, which has not been feasible with existing approaches. To directly test this possibility, we established a high-throughput workflow combining time-lapse imaging of dual-fluorescent HSV-1 reporter viruses with automated single-cell image analysis (Fig. 3A). In these viruses, TagRFP and Venus were fused to IE proteins and to either an E or an L protein, enabling direct visualization of the temporal cascade of viral gene expression. For long-term single-cell tracking, the accuracy of trajectory assignment must be verifiable to ensure reliable downstream analysis. We therefore employed a “sparse-labeling” strategy, in which 10% of cells were pre-labeled with CellTrace Far Red and mixed with unlabeled cells, thereby preserving the dense culture environment. This configuration enabled robust segmentation using a deep-learning–based algorithm (Cellpose3) ^24^ and allowed visual confirmation of tracking accuracy by following each labeled cell unambiguously across frames (S-Movie 1). Cascade dynamics were quantified using an “onset” metric, defined for each individual cell as the first time point at which fluorescence exceeded a threshold relative to that cell’s own baseline (Fig. 3A). We confirmed that CellTrace labeling did not affect TagRFP or Venus fluorescence accumulation at the population level (S-Fig. 3A and B) and that imaging-based measurements closely matched conventional flow cytometry (S-Fig. 3C and D), demonstrating that our imaging system achieves quantitative accuracy comparable to flow cytometry and enables longitudinal single-cell analysis of HSV-1 gene-expression dynamics in rICP47/vUs11-infected wild-type, CALM1-KO, and CALM1/2-KO cells.

**Fig. 3.**
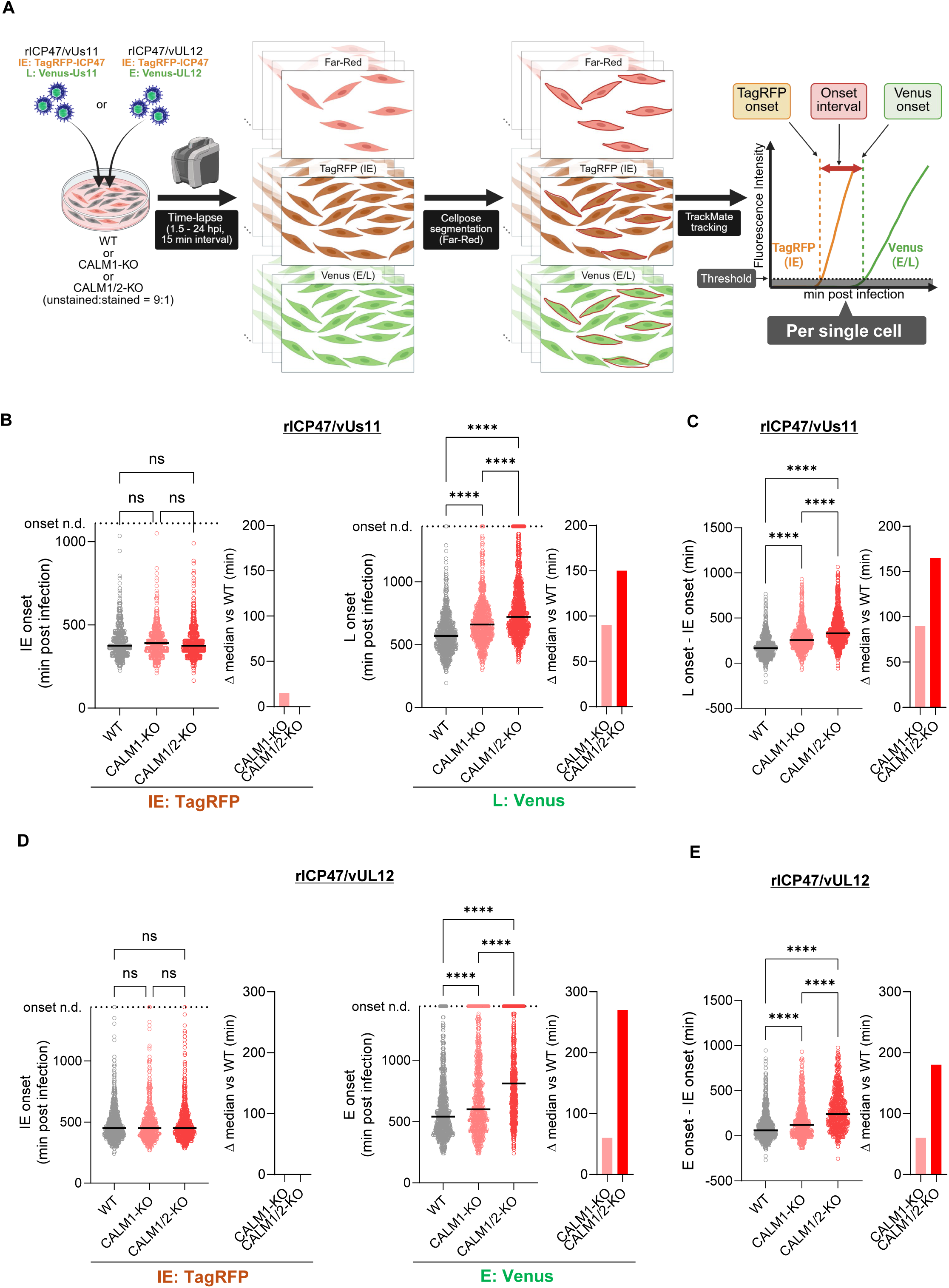
CaM deficiency delays the onset of E/L gene expression in individual cells. (A) Schematic workflow of fluorescence onset analysis. Indicated cells were infected with dual-fluorescent reporter HSV-1 (rICP47/vUs11 or rICP47/vUL12), in which 10% of the cell population had been pre-labeled with CellTrace Far Red. Time-lapse imaging was performed every 15 min from 1.5 to 24 h post-infection. Far Red–labeled cells were segmented using Cellpose and tracked using TrackMate to obtain single-cell trajectories of TagRFP and Venus fluorescence intensities. For each tracked cell, the threshold was defined as the baseline fluorescence (mean of the first five time points) plus five times the baseline standard deviation. The first time point at which the fluorescence intensity exceeded this threshold for five consecutive frames was defined as the onset of TagRFP (IE) or Venus (E or L) expression. (B and C) HeLa WT, CALM1-KO, and CALM1/2-KO cells were infected with rICP47/vUs11 at an MOI of 5 and imaged every 15 min from 1.5 to 24 h post-infection. For each cell, the onsets of TagRFP (IE) and Venus (L) fluorescence were determined from time-lapse trajectories. (B) Dot plots showing IE onset (left) and L onset (right) times (min post-infection) for individual cells. Cells in which onset was not detected are plotted above the dashed line (onset n.d.). (C) Time interval between IE and L onset (L onset − IE onset) in cells where both onsets were defined. (D and E) HeLa WT, CALM1-KO, and CALM1/2-KO cells were infected with rICP47/vUL12 at an MOI of 5 and imaged under the same conditions as in (B and C). (D) Dot plots showing IE onset (left) and E onset (right) times (min post-infection) for individual cells. Cells in which onset was not detected are plotted above the dashed line (onset n.d.). (E) Time interval between IE and E onset in cells where both onsets were defined. In (B to E), each dot represents one cell and bars represent medians. Adjacent bar graphs indicate the differences in median values between WT and each KO population. Cells in which onset was not detected within the imaging period are plotted at the maximum observed onset value plus 60 min (above the dashed line, onset n.d.) and were included in statistical analyses with this imputed value. The number of analyzed cells was as follows: (B) WT, n = 842; CALM1-KO, n = 886; CALM1/2-KO, n = 1056. (C) WT, n = 839; CALM1-KO, n = 880; CALM1/2-KO, n = 1011. (D) WT, n = 877; CALM1-KO, n = 704; CALM1/2-KO, n = 798. (E) WT, n = 851; CALM1-KO, n = 652; CALM1/2-KO, n = 662. Data represent pooled measurements from two independent experiments. Statistical analyses were performed using the Kruskal–Wallis test followed by Dunn’s multiple-comparison test; ****, p < 0.0001; ns, not significant.

Using rICP47/vUs11, we first monitored progression from IE to L gene expression at the single-cell level. The onset of IE (TagRFP) protein expression was comparable across cell types, whereas the onset of L (Venus) protein expression was stepwise delayed with increasing numbers of disrupted CALM genes (Fig. 3B), consistent with the preferential reduction of E and L mRNA abundance observed by qPCR at the population level (Fig. 2A and B). Accordingly, the IE–L onset interval was incrementally prolonged among cells in which both onsets could be defined (Fig. 3C), indicating that CaM deficiency delays the transition from IE to downstream gene expression at the single-cell level rather than affecting the initiation of IE expression itself.

To evaluate E gene expression, we constructed rICP47/vUL12, in which TagRFP and Venus were fused to IE protein ICP47 and the E protein UL12, respectively (S-Fig. 4A and B). This reporter virus grew comparably to wild-type virus (S-Fig. 4C and D), similar to rICP47/vUs11 ^22^. The onset of IE (TagRFP) protein expression at the single-cell level was comparable across cell types, whereas both the onset of E (Venus) protein expression and the IE–E onset interval were stepwise delayed with increasing numbers of disrupted CALM genes (Fig. 3D and E). Notably, the fraction of cells in which E or L onset was not detected by 24 h post-infection increased stepwise with the number of disrupted CALM genes (Fig. 3B, D, S–Fig. 5A and B), suggesting that such progressive onset delays can, in extreme cases, prevent cells from initiating E and L gene expression within the timeframe of a productive infection cycle. Delays in E and L onsets, as well as prolonged IE–E/L intervals, were restored in CALM1-KO cells complemented with Flag-CaM (S-Fig. 5C–F).

Because the delayed rise in E and L protein signals could reflect either delayed onset or slower post-onset protein accumulation, we next sought to distinguish between these possibilities. We analyzed the impact of CaM deficiency on the temporal accumulation profiles of IE, E, and L proteins using single-cell fluorescence trajectories (Fig. 4A). In rICP47/vUs11-infected cells, the population-averaged TagRFP fluorescence trajectory (reflecting IE protein expression) (Fig. 4A, left) was similar among wild-type, CALM1-KO, and CALM1/2-KO cells, whereas the Venus fluorescence trajectory (reflecting L protein expression) exhibited a delayed rise in CaM-deficient cells (Fig. 4B), consistent with the delayed onset of L protein expression (Fig. 3B). To isolate post-onset accumulation kinetics from the effect of variable onset timing, we aligned individual cell trajectories to their respective IE (TagRFP) or L (Venus) onset and averaged fluorescence intensity at each time point relative to onset (Fig. 4A, right, C and D). The resulting accumulation profiles of both IE (TagRFP) and L (Venus) fluorescence were largely comparable among wild-type, CALM1-KO, and CALM1/2-KO cells. Similar results were obtained for the E gene reporter rICP47/vUL12 (Fig. 4E to G).

**Fig. 4.**
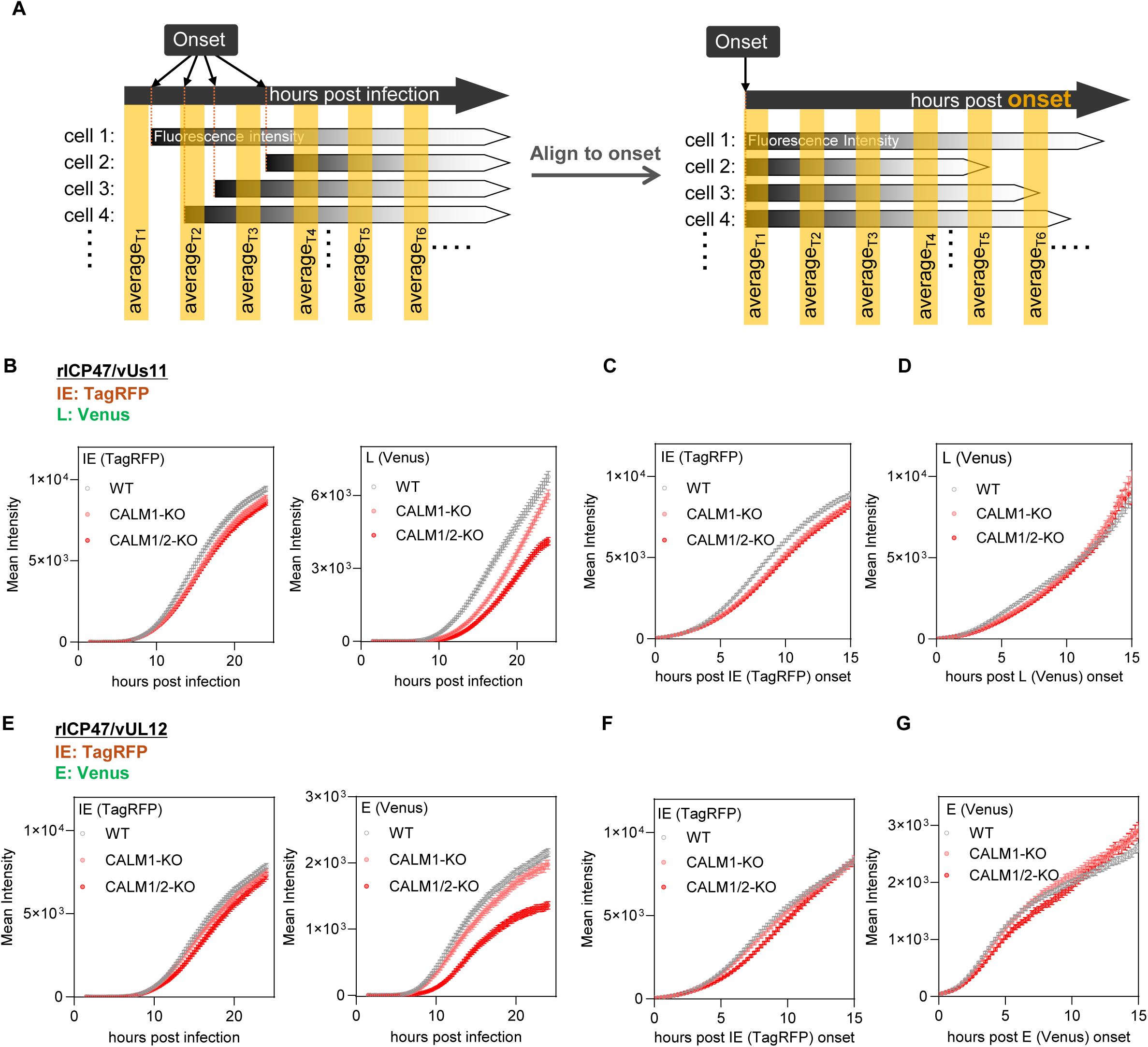
Single-cell temporal alignment analysis of HSV-1 protein accumulation kinetics. (A) Schematic of the temporal alignment approach. Left: single-cell fluorescence trajectories plotted against hours post-infection, with variable onset times across cells. Right: the same trajectories aligned by setting each cell’s onset as time 0, enabling comparison of post-onset accumulation kinetics. (B to D) Population-averaged fluorescence trajectories from rICP47/vUs11 time-lapse imaging, reanalyzed from the dataset shown in Fig. 3B. (B) Population-averaged mean fluorescence intensity of IE (TagRFP) and L (Venus) proteins plotted against hours post-infection. (C and D) The single-cell fluorescence trajectories used for (B) were computationally aligned by setting the onset of IE (TagRFP) (C) or L (Venus) (D) as time 0 for each cell. Population-averaged mean fluorescence intensity is plotted against hours post-onset. The number of cells analyzed ranged from n = 839 to 780 (WT), n = 880 to 736 (CALM1-KO), and n = 1011 to 952 (CALM1/2-KO) in (C); and from n = 839 to 389 (WT), n = 880 to 193 (CALM1-KO), and n = 1011 to 114 (CALM1/2-KO) in (D). (E to G) Population-averaged fluorescence trajectories from rICP47/vUL12 time-lapse imaging, reanalyzed from the dataset shown in Fig. 3D. (E) Population-averaged mean fluorescence intensity of IE (TagRFP) and E (Venus) proteins plotted against hours post-infection. (F and G) The single-cell fluorescence trajectories used for (E) were computationally aligned by setting the onset of IE (TagRFP) (F) or E (Venus) (G) as time 0 for each cell. Population-averaged mean fluorescence intensity is plotted against hours post-onset. The number of cells analyzed ranged from n = 851 to 660 (WT), n = 652 to 512 (CALM1-KO), and n = 662 to 540 (CALM1/2-KO) in (F); and from n = 851 to 461 (WT), n = 652 to 284 (CALM1-KO), and n = 662 to 161 (CALM1/2-KO) in (G). In (C, D, F, and G), the number of cells analyzed decreases over time as individual trajectories end at different durations after alignment. Data are presented as mean ± SE.

Together, these results demonstrate that, at the single-cell level, CaM regulates the onset timing of E and L protein expression, whereas their post-onset accumulation profiles are largely preserved under CaM deficiency, establishing CaM as a regulator of the tempo of the HSV-1 gene-expression cascade.

### CaM deficiency reduces progeny virus production by reshaping the distribution of viral gene-expression outcomes across individual cells

In time-lapse imaging of rICP47/vUs11-infected CALM1-KO and CALM1/2-KO cell populations, the Venus signal—serving as a proxy for L protein expression—showed a delayed rise during the early phase of infection before gradually reaching levels comparable to those in wild-type cells (Fig. 4B and S–Fig. 6). In line with the imaging results, flow cytometric analysis at 12 h post-infection showed that the fraction of Venus-positive cells was reduced in CaM-deficient cells (1.3-fold in CALM1-KO cells and 2.0-fold in CALM1/2-KO cells) relative to wild-type cells, whereas the fraction of TagRFP-positive cells, representing IE protein expression, was comparable across cell types (S-Fig. 7A to C). Furthermore, the delayed rise in the Venus signal (S-Fig. 6) closely resembled the kinetics of viral replication observed under the same MOI (Fig. 2C), suggesting a potential link between delayed L protein accumulation and reduced progeny virus production. However, our previous analysis using the same reporter virus showed that the relationship between L protein accumulation and progeny virus production is threshold-dependent at the subpopulation level ^22^. Thus, progeny virus production initiates only when Venus fluorescence exceeds a defined threshold in individual cells, indicating that a certain level of L protein accumulation is required for producing infectious virions ^22^. These observations raise the possibility that the reduced progeny virus yield observed in CaM-deficient cells results from a decreased proportion of virus-producing cells within the population, owing to delayed L protein expression that postpones the time at which individual cells reach the expression threshold required for productive infection. Consistent with this possibility, the fraction of Venus^high^ cells—defined as the virus-producing population based on a Venus fluorescence threshold established in our previous study ^22^ —was markedly reduced by 2.4-fold in CALM1-KO cells and 7.4-fold in CALM1/2-KO cells relative to wild-type cells (Fig. 5A and B), with the magnitude of the reduction substantially exceeding that observed for the Venus-positive population (S-Fig. 7C). These results suggest that CaM deficiency reshapes the distribution of infection outcomes across individual cells by reducing the fraction of cells that accumulate sufficient L protein for progeny virus production.

**Fig. 5.**
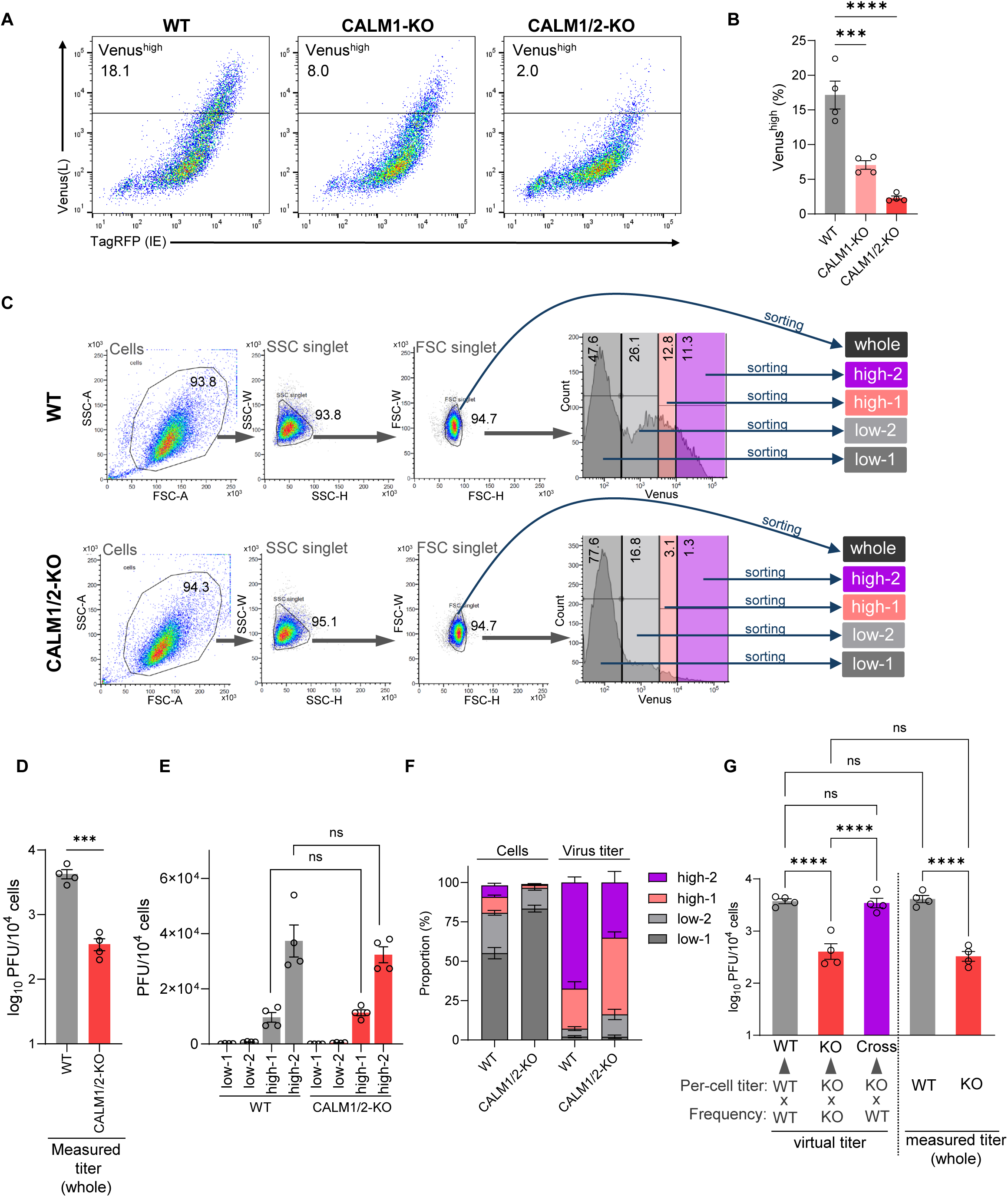
CaM deficiency reduces progeny virus production by altering the heterogeneity of viral gene expression. (A and B) HeLa WT, CALM1-KO, and CALM1/2-KO cells were infected with rICP47/vUs11 at an MOI of 5. At 12 h post-infection, cells were analyzed by flow cytometry. (A) Representative flow cytometry plots showing TagRFP (IE) and Venus (L) fluorescence. The gate for the Venus^high^ population (virus-producing cells) is shown, with the percentage of cells within this gate indicated. (B) Quantification of the percentage of Venus^high^ cells from (A). (C to G) HeLa WT and CALM1/2-KO cells were infected as in (A and B) and subjected to fluorescence-activated cell sorting (FACS) at 12 h post-infection. (C) Gating strategy for cell sorting. Single cells (SSC- and FSC singlet) were sorted into the whole population or fractionated into four subpopulations (low-1, low-2, high-1, and high-2) based on Venus (L) fluorescence intensity. (D) Viral titers (PFU per 10⁴ cells) from the sorted “whole” singlet population. (E) Viral titers of each of the four sorted subpopulations in (C) (i.e., the per-cell titers used in G). (F) Stacked bar graphs showing, for WT and CALM1/2-KO (KO) cells, the subpopulation composition (“Cells”) and each subpopulation’s contribution to the total viral yield (“Virus titer”), both expressed as percentages. (G) Virtual titer analysis. For each Venus-defined subpopulation, the product of its frequency among infected cells (“Frequency”) and its per-cell titer (“Per-cell titer”) was calculated, and these products were summed across subpopulations to obtain the virtual titer. The genotype source of each parameter (WT or CALM1/2-KO) is indicated below each bar: the “WT” and “KO” bars use the Frequency and Per-cell titer from the same genotype, whereas the “Cross” bar combines the WT Frequency with the CALM1/2-KO Per-cell titer. Measured whole-population titers (as in D) are shown on the right for comparison. Each value represents the mean ± SE of four (B, D, E, F, and G) biological replicates. Statistical analyses were performed using one-way ANOVA followed by Tukey’s multiple-comparison test (B, E, and G), or a Welch’s *t*-test (D). ***, p < 0.001; ****, p < 0.0001; ns, not significant.

**Fig. 6.**
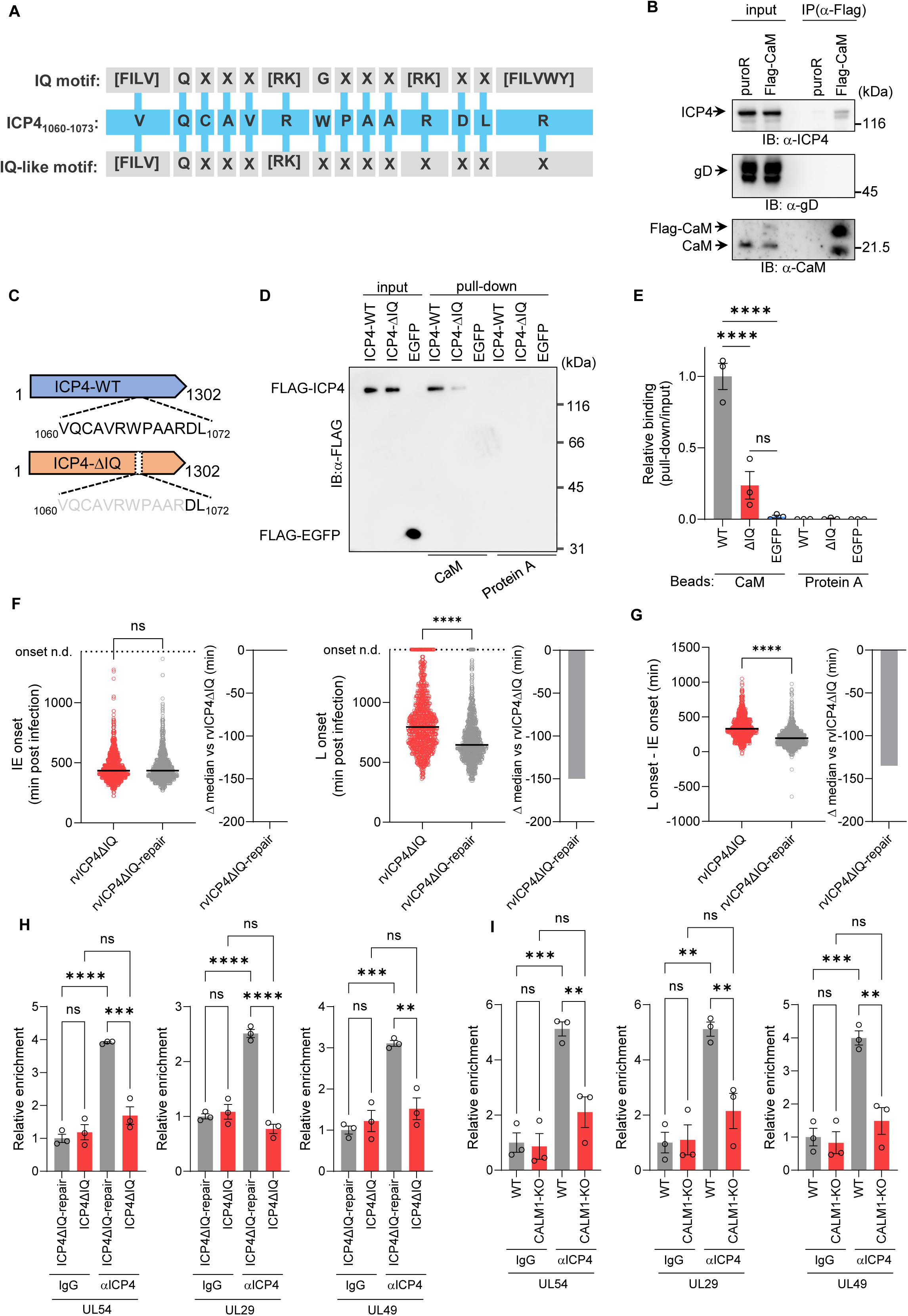
CaM promotes ICP4 binding to the viral genome through its IQ-like motif to regulate HSV-1 gene expression and replication. (A) Alignment of the amino acid sequence of HSV-1 ICP4 (aa 1060–1073) with the canonical IQ motif and the IQ-like motif. (B) HeLa/puroR or HeLa/Flag-CaM cells were infected with wild-type HSV-1(F) at an MOI of 5. At 9 h post-infection, cell lysates were immunoprecipitated (IP) with an anti-Flag antibody. Input lysates and immunoprecipitates were analyzed by immunoblotting with the indicated antibodies. (C) Schematic diagram of the structure of wild-type ICP4 (ICP4-WT) and the IQ-like motif deletion mutant (ICP4-ΔIQ), which lacks amino acids 1060–1070. (D and E) Purified Flag-tagged ICP4-WT, ICP4-ΔIQ, or EGFP were incubated with calmodulin (CaM) Sepharose beads or Protein A Sepharose (control) beads. Bound proteins were detected by immunoblotting with an anti-Flag antibody (D). (E) Quantification of the relative binding intensity (pull-down/input) from (D). (F and G) HeLa cells were infected with rICP47/vUs11/ICP4ΔIQ (rvICP4ΔIQ) or its repaired virus (rvICP4ΔIQ-repair) at an MOI of 5 and analyzed by time-lapse imaging as in Fig. 3A. (F) Dot plots showing IE onset (TagRFP, left) and L onset (Venus, right) times (min post-infection) for individual cells. Cells in which onset was not detected are plotted above the dashed line (onset n.d.) with an imputed value (maximum observed onset + 60 min) and were included in statistical analyses. (G) Time interval between IE and L onset (L onset − IE onset) in cells where both onsets were defined. (H) HeLa cells were infected with ICP4ΔIQ or ICP4ΔIQ-repair virus at an MOI of 5 and fixed at 3.5 h post-infection. ChIP-qPCR was performed using an anti-ICP4 antibody or control IgG to quantify ICP4 occupancy at the 5′ regions of UL54 (IE), UL29 (E), and UL49 (L). (I) WT or CALM1-KO HeLa cells were infected with wild-type HSV-1(F) at an MOI of 5 and analyzed by ChIP-qPCR at 3.5 h post-infection as in (H). Data are representative of three independent experiments (B and D). Each value represents the mean ± SE of three (E, H, and I) biological replicates. In (F and G), each dot represents one cell and bars indicate medians. Adjacent bar graphs indicate the differences in median values relative to rvICP4ΔIQ. The number of analyzed cells was as follows: (F) rvICP4ΔIQ, n = 956; rvICP4ΔIQ-repair, n = 1079; (G) rvICP4ΔIQ, n = 912; rvICP4ΔIQ-repair, n = 1065. Data represent pooled measurements from two independent experiments. Statistical analyses were performed using one-way ANOVA followed by Tukey’s multiple-comparison test (E, H, and I) or Mann–Whitney U-test (F and G). **, p < 0.01; ***, p < 0.001; ****, p < 0.0001; ns, not significant.

To directly test whether the reduced fraction of Venus^high^ cells account for the decreased progeny virus production in CaM-deficient cells, wild-type and CALM1/2-KO cells infected with rICP47/vUs11 at an MOI of 5 were sorted at 12 h post-infection, a time point at which viral yield was reduced under the same MOI (Fig. 2C), into the whole singlet population and four subpopulations defined by Venus intensity (low-1, low-2, high-1, and high-2 subpopulations) (Fig. 5C). The Low-1 and low-2 subpopulations corresponded to cells that barely produce detectable progeny virus, whereas the high-1 and high-2 subpopulations represented the major virus-producing cells, with high-2 approaching the virus production plateau, as defined previously ^22^. Consistent with Fig. 2C, virus titers from whole sorted populations were significantly lower in CALM1/2-KO cells than in wild-type cells (Fig. 5D). In contrast, titers per 10⁴ cells within the high-1 and high-2 subpopulations were comparable between cell types, indicating that CaM deficiency does not affect progeny virus production on a per-cell basis. Instead, the combined frequency of the high-1 and high-2 subpopulations decreased from 17.4% in wild-type cells to 2.5% in CALM1/2-KO cells, accounting for more than 80% of the progeny virus yield in both cell types (Fig. 5E and F). These results indicate that the overall virus yield defect largely reflects a reduced abundance of virus-producing cells. To quantitatively test whether changes in subpopulation distribution alone account for the yield defect, we calculated “virtual titers” by multiplying the frequency of each subpopulation by its per-cell titer and summing these values across subpopulations. Virtual titers closely matched the measured titers of the whole population for both wild-type and CALM1/2-KO cells, validating this approach (Fig. 5G). We then performed a cross-combination analysis in which subpopulation frequencies from one genotype were combined with per-cell titers from the other. When wild-type subpopulation frequencies were combined with CALM1/2-KO per-cell titers, the resulting virtual titers were significantly higher than those calculated using only CALM1/2-KO-derived values and approached wild-type levels (Fig. 5G), indicating that the shift in subpopulation composition is sufficient to account for the yield defect in CaM-deficient cells. Similar results were obtained with CALM1-KO cells (S-Fig. 7D to H). In CALM1-KO cells expressing Flag-CaM, both the fraction of Venus^high^ subpopulations and the viral yield of the whole population were increased relative to CALM1-KO cells (S-Fig. 8A to D), whereas per-cell titers of high-1 and high-2 subpopulations remained comparable (S-Fig. 8E). Cross-combination analysis confirmed that the recovery of viral yield reflects restoration of subpopulation distribution (S-Fig. 8F and G).

Taken together with the time-lapse imaging results (Fig. 3 and S–Fig. 5), these findings indicate that CaM-dependent regulation of the tempo of the viral gene-expression cascade at the single-cell level determines the fraction of cells that exceed the L protein expression threshold required for virus production, thereby governing progeny virus production at the population level.

### CaM promotes HSV-1 gene expression and replication by facilitating ICP4 binding to viral genome through its IQ-like motif

Because CaM deficiency preferentially suppressed E and L gene expression, we reasoned that CaM might interact with a specific IE protein and affect its ability to activate downstream gene expression. To explore this possibility, we queried the Calmodulin Target Database ^25^ for consensus calmodulin-binding motifs within HSV-1 IE proteins. Among the IE proteins, only ICP4 contained a predicted consensus sequence, harboring an IQ-like motif. Specifically, amino acids 1060–1073 of ICP4 closely resemble the canonical IQ consensus and matched the IQ-like motif (Fig. 6A) ^26^. Immunoprecipitation of Flag-tagged calmodulin from wild-type HSV-1(F)-infected HeLa/Flag-CaM cells resulted in co-precipitation of ICP4, indicating that CaM associates with ICP4 in HSV-1-infected cells (Fig. 6B). To assess direct binding via the IQ-like motif, we purified Flag-tagged ICP4 and a mutant lacking amino acids 1060–1070 of this motif (ICP4ΔIQ) (Fig. 6C and S–Fig. 9A) and tested their interaction with calmodulin in pull-down assays. Flag-ICP4 bound to calmodulin-conjugated beads, whereas ICP4ΔIQ displayed markedly reduced binding (Fig. 6D and E). This interaction was specific, as EGFP showed no binding to calmodulin-conjugated beads and Protein A beads did not pull down ICP4. Together, these results indicate that calmodulin interacts with ICP4 primarily through its IQ-like motif.

To assess the functional role of the IQ-like motif during viral infection, we generated recombinant viruses carrying the ΔIQ mutation in ICP4 (ICP4ΔIQ) and its repaired virus (ICP4ΔIQ-repair) (S-Fig. 4A). The ΔIQ mutation did not alter ICP4 protein expression in HSV-1-infected cells (S-Fig. 9B). However, at both MOIs of 5 and 0.05, ICP4ΔIQ yielded significantly lower titers than wild-type HSV-1(F) or ICP4ΔIQ-repair (S-Fig. 9C and D). qPCR analysis showed reduced mRNA levels of all viral genes examined in ICP4ΔIQ-infected cells compared with wild-type HSV-1(F) and ICP4ΔIQ-repair (S-Fig. 9E). The reduction was modest for the IE genes ICP0 and ICP27, but more pronounced for the E genes ICP8 and UL50 and the L genes UL49 and Us2, recapitulating the expression pattern in CALM1-KO and CALM1/2-KO cells.

To further investigate the role of the ICP4 IQ-like motif in HSV-1 gene expression dynamics, we introduced the ΔIQ mutation into rICP47/vUs11 to generate rICP47/vUs11/ICP4ΔIQ (rvICP4ΔIQ) and its repaired counterpart, rICP47/vUs11/ICP4ΔIQ-repair (rvICP4ΔIQ-repair) (S-Fig. 4A). Time-lapse imaging indicated that the mutation did not affect IE onset (TagRFP) but delayed the onset of L (Venus) expression and prolonged the IE–L interval (Fig. 6F and G). Flow cytometry showed that the frequency of Venus^high^ cells was reduced in rvICP4ΔIQ-infected populations compared with the repaired virus (S-Fig. 10A and B). To quantitatively assess whether this reduced proportion of virus-producing cells accounts for the yield defect caused by the ΔIQ mutation, we sorted cells into subpopulations based on Venus intensity and measured titers in each (S-Fig. 10C to E). The ΔIQ mutation significantly reduced titers in the whole population but did not significantly affect titers within the virus-producing high-1 and high-2 subpopulations (S-Fig. 10D and E). In rvICP4ΔIQ-infected cells, high-1 and high-2 cells, which represented only 2.2% of the total population, contributed 87.5% of the viral output (S-Fig. 10F). Cross-combination analysis demonstrated that the yield defect resulted from a reduced proportion of these subpopulations (S-Fig. 10G).

ICP4 binds to the HSV-1 genome in infected cells and activates the expression of E and L genes by recruiting cellular general transcription factors to viral promoters ^27–31^. The finding that deletion of the IQ-like motif preferentially attenuated E and L gene expression suggested that this mutation might impair the DNA-binding activity of ICP4. To examine this, we performed ChIP–qPCR to compare ICP4 occupancy at the 5′ regions of the IE gene UL54, the E gene UL29, and the L gene UL49 in cells infected with either ICP4ΔIQ or ICP4ΔIQ-repair viruses. The ΔIQ mutation was associated with reduced ICP4 binding at all of these regions (Fig. 6H). This reduction was recapitulated in CALM1-KO cells infected with wild-type HSV-1(F) (Fig. 6I). Under these conditions, total ICP4 protein levels were comparable (S-Fig. 10H and I). Taken together, these results indicate that CaM promotes ICP4 binding to viral promoters through its IQ-like motif, thereby facilitating timely activation of E and L gene expression following IE expression in individual cells and ensuring efficient HSV-1 replication at the population level.

### Pharmacological inhibition of CaM protects mice from lethal HSV-1 infection

Because our in vitro data suggested that CaM facilitates HSV-1 replication by regulating the tempo of viral gene expression, we next examined whether pharmacological inhibition of CaM could influence HSV-1 pathogenesis in vivo. Administration of the CaM inhibitor W-7 protected mice from lethal HSV-1 infection following intracranial inoculation (Fig. 7A). Of note, brain viral titers at 2–3 days post-infection were reduced to below the detection limit in most animals (Fig. 7B). The protective effect of W-7 against HSV-1–induced mortality was also greater than that observed with the standard antiviral drug acyclovir (ACV) (Fig. 7C). Together, these results identify CaM as a potential therapeutic target in HSV-1 encephalitis.

**Fig. 7.**
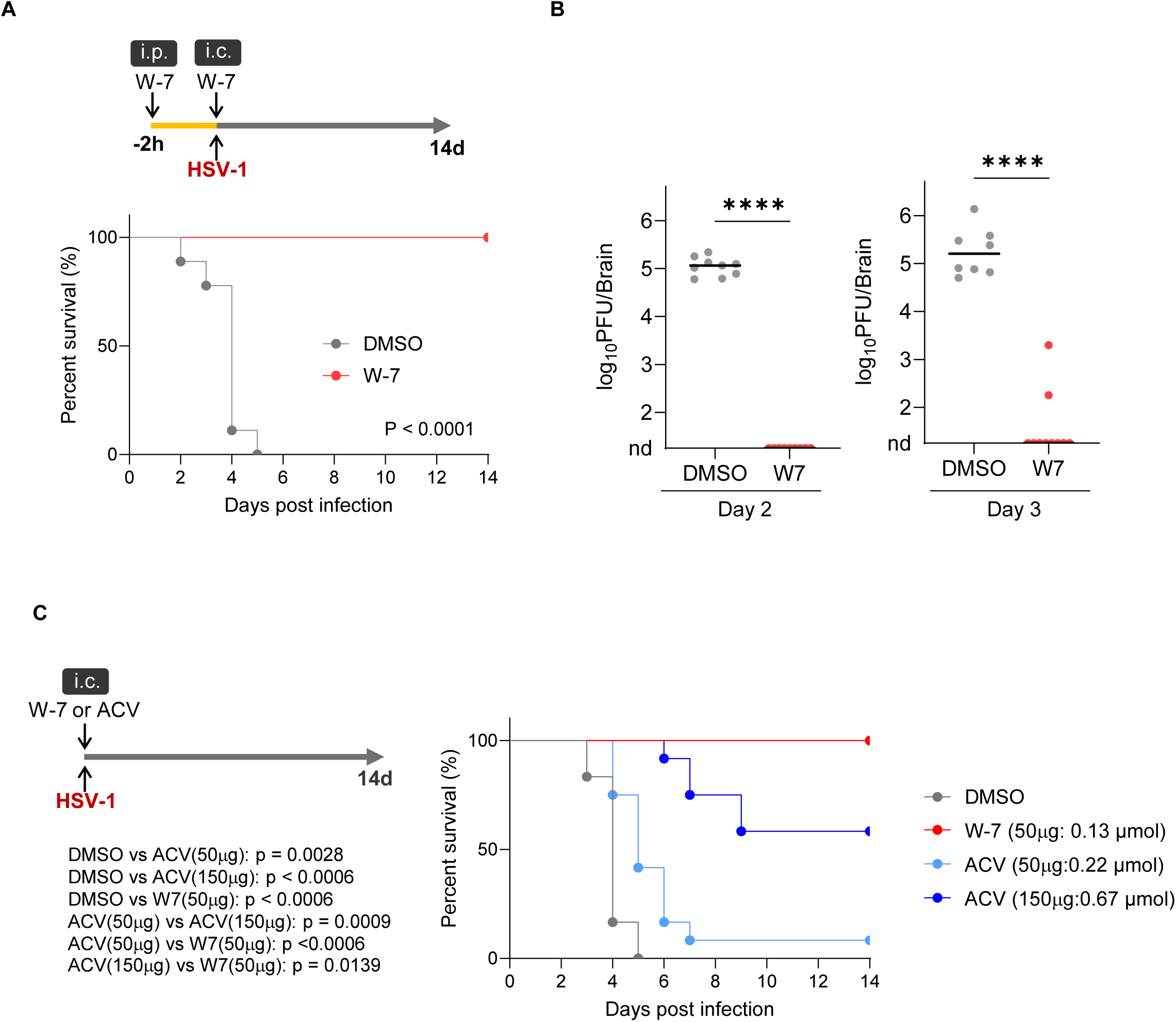
Pharmacological inhibition of calmodulin protects mice from lethal HSV-1 infection. (A) Schematic of W-7 treatment and survival curve. Three-week-old female ICR mice (n=9 per group) were administered W-7 (i.p. 125 µg at 2 h before infection) and subsequently infected intracranially (i.c.) with wild-type HSV-1(F) (5 × 10² PFU) mixed with W-7 (50 µg) or DMSO (vehicle control). Survival was monitored for 14 days. The P value was calculated using the log-rank test. (B) Viral titers in the brain at 2 and 3 days post-infection (dpi) from mice treated as in (A). Day 2: DMSO (n=9), W-7 (n=9); Day 3: DMSO (n=8), W-7 (n=10). Samples with non-detectable virus are shown at the “nd” position on the graph. For statistical analysis, these non-detectable samples were imputed with a value of 18 PFU/brain, which is the highest value below the detection limit. P values were calculated using the Mann–Whitney U-test. ****, p < 0.0001. (C) Schematic of W-7 or acyclovir (ACV) treatment and survival curve. Mice (n=12 per group) were infected i.c. with wild-type HSV-1(F) mixed with DMSO (vehicle control), W-7 (50 µg: 0.13 μmol), ACV (50 µg: 0.22 μmol), or ACV (150 µg: 0.67 μmol). Survival was monitored for 14 days. P values shown were calculated using the log-rank test with Holm’s correction for multiple comparisons.

## DISCUSSION

Viral replication exhibits remarkable heterogeneity among individual infected cells, ^2–6^ and in many DNA viruses, temporally ordered cascades of gene expression ^1^ are likely a major source of this variability. However, how the tempo of cascade progression is regulated at the single-cell level and how this variability shapes population-level replication outcomes have remained unclear, in part due to the lack of experimental approaches capable of resolving gene-expression dynamics in individual cells. In this study, we establish a high-throughput live-cell imaging platform that enables single-cell tracking of the temporal progression of the HSV-1 gene-expression cascade. By integrating this imaging platform with cell sorting–based analyses linking single-cell gene expression to population-level virus production, together with single-cell RNA sequencing, we demonstrate that the tempo of cascade progression is indeed regulated at the single-cell level and identify CaM as a critical host factor controlling this process. A key insight of this study is that CaM regulates population-level viral replication by controlling the tempo of the viral gene-expression cascade—specifically, by advancing the onset of E and L gene expression following IE expression in individual cells. Our findings reveal a previously unrecognized regulatory layer embedded within the canonical gene-expression cascade—the control of its tempo. Importantly, this temporal regulation at the single-cell level reshapes the distribution of infection outcomes by increasing the fraction of cells that exceed the threshold of viral gene expression required for progeny virus production without altering their burst size, thereby promoting progeny virus production at the population level. Thus, our study establishes a direct mechanistic link between single-cell gene-expression dynamics and population-level viral output, demonstrating that population-level viral replication is governed not by the average behavior of individual cells but by the distribution of heterogeneous cellular states within the population. Furthermore, our integrated experimental and analytical framework provides a broadly applicable approach to dissect how temporal dynamics, including single-cell heterogeneity, are translated into population-level outcomes.

Our scRNA-seq analysis revealed marked cell-to-cell variability in CaM mRNA abundance among host cells. This finding is consistent with a previous single-cell proteomic study showing substantial heterogeneity in CaM protein abundance across individual mammalian cells ^32^. Such variability likely reflects differences in host cell states, including signaling activity and stress responses ^21^. Given that heterogeneity in viral infection is closely associated with progression through the viral gene-expression cascade ^5,33^, and that CaM regulates the tempo of the HSV-1 gene-expression cascade at the single-cell level, our results suggest that variability in cellular CaM abundance may influence the tempo of cascade progression, thereby contributing to infection heterogeneity. Consistent with this possibility, stepwise reduction of CaM expression in CALM1-KO and CALM1/2-KO cells was accompanied by a corresponding stepwise delay in progression from IE to E and L phases at the single-cell level, supporting a causal link between CaM abundance, cascade tempo, and infection outcomes. We propose that HSV-1 may exploit this intrinsic variability as a survival strategy, diversifying the timing of progeny virus production without altering the final progeny yield (burst size) in individual cells. This temporal diversification is consistent with a “bet-hedging” strategy in which asynchronous progression of infection reduces the risk of simultaneous immune detection and clearance ^34^. In this model, temporal dispersion of viral gene expression and progeny production across infected cells enhances viral persistence under fluctuating host conditions. Similar strategies have been described in other viral systems, including the lytic–lysogenic switch of bacteriophages and latency establishment by HIV ^35–37^, suggesting that this principle may be broadly applicable.

Mechanistically, we identify ICP4 as a key mediator of CaM-dependent temporal control. ICP4 is the major transcriptional activator of HSV-1 ^12,17^, and we show that it interacts with CaM through an IQ-like motif. Disruption of this interaction, either by deletion of the IQ-like motif or by CaM deficiency, reduces ICP4 occupancy at viral promoters and delays progression of the gene-expression cascade at the single-cell level. These findings support a model in which CaM enhances ICP4 function to ensure timely activation of E and L genes in individual cells. Thus, the CaM–ICP4 axis acts as a molecular timer that governs the pace of viral transcriptional progression.

Finally, we demonstrate that pharmacological inhibition of CaM by W-7 markedly protects mice from lethal HSV-1 infection in vivo. This effect exceeded that of acyclovir (ACV), a conventional antiviral agent, under the conditions tested. Whereas conventional antiviral agents target viral enzymes, CaM inhibition disrupts host-dependent regulation of viral gene-expression dynamics, thereby reducing the fraction of cells that progress to productive infection. These findings suggest a conceptually distinct antiviral strategy and identify CaM as a promising host-directed therapeutic target. However, given the pleiotropic roles of CaM, further studies will be required to evaluate the specificity and safety of this approach.

## MATERIALS AND METHODS

### Cells and viruses

HeLa, Plat-GP, and Vero cells were described previously ^38,39^. Wild-type HSV-1(F), YK410 (rICP47/vUs11), and YK651 (vUL12) were described previously ^22,40,41^.

### Generation of a recombinant virus

Recombinant virus YK414 (rICP47), in which ICP47 was tagged with TagRFP, was generated by two-step Red-mediated mutagenesis in *Escherichia coli* GS1783 containing pYEbac102Cre, a full-length infectious wild-type HSV-1(F) clone, as described previously^40,42^, using the template plasmids and primers listed in Table S1. Recombinant viruses YK416 (ICP4ΔIQ) and YK418 (rICP47/vUs11/ICP4ΔIQ) were generated by two rounds of two-step Red-mediated mutagenesis, as described previously ^40,42^, except that the *E. coli* clone, template plasmids, and primers listed in Table S1 were used. Recombinant viruses YK417 (ICP4ΔIQ-repair) and YK419 (rICP47/vUs11/ICP4ΔIQ-repair) were generated by one round of two-step Red-mediated mutagenesis, as described previously ^40,42^, except that the E. coli clone, template plasmids, and primers listed in Table S1 were used to restore a single copy of ICP4. The resulting BAC DNAs were prepared from *E. coli* and electroporated into Vero cells to reconstitute infectious HSV-1. The reconstituted viruses were subjected to three rounds of plaque purification to obtain clonal stocks, whose sequences were verified to confirm restoration of the ICP4 mutation. YK415 (rICP47/vUL12), in which ICP47 and UL12 were tagged with TagRFP and mVenus, respectively, were generated by co-infecting Vero cells with YK414 (rICP47) and YK651 (vUL12), followed by three rounds of plaque purification of progeny viruses exhibiting both Venus and TagRFP fluorescence. Correct insertion of each fluorescent tag was subsequently confirmed by sequencing across the insertion site.

### Plasmids

Plasmids pFLAG-EGFP, pFLAG-ICP4, pFLAG-UL29, pFLAG-ICP0, pFLAG-UL54, pFLAG-Us2, pFLAG-UL49, and pFLAG-UL50 were described previously ^39^. Plasmids constructed in this study were generated by standard PCR-based cloning or oligonucleotide insertion into the following recipient vectors: p-FLAG-CMV2 (Sigma) for pFLAG-ICP4Δ1060-1070 and pFLAG-18s, pMx-puro ^43^ for pMx-puro-FlagCaM and pMx-puro-FlagCaMv2, px459 ^44^ for px459-CALM1, and px458 ^44^ for px458-CALM2. Details of plasmid construction, including PCR templates and oligonucleotide sequences, are provided in Table S2.

### Single-cell RNA sequencing (scRNA-seq) and data analysis

#### scRNA-seq

HeLa cells were mock infected or infected with wild-type HSV-1(F) at a multiplicity of infection (MOI) of 5. At 13 h post-infection, cells were detached with trypsin and used for single-cell RNA-Seq analysis on the C1 system (Fluidigm, South San Francisco, CA, USA) with the C1 Single-Cell Auto Prep Reagent Kit for mRNA Seq (Fluidigm) and the C1 Single-Cell Auto Prep IFC for mRNA Seq (Fluidigm). RNA-seq libraries were constructed according to the manufacturers’ protocols. Briefly, single cells were captured on the C1 IFC and partitioned into 96 independent reaction chambers. First-strand cDNA synthesis and whole-transcriptome amplification were performed using the SMART-Seq v4 Ultra Low Input RNA Kit for the Fluidigm C1 System (Takara-Clontech, Mountain View, CA, USA). Sequencing libraries were then prepared with the Nextera XT DNA Sample Preparation kit (Illumina, San Diego, CA, USA), assessed for quality and quantity using a Bioanalyzer 2100 (Agilent Technologies, Santa Clara, CA, USA), and sequenced on an Illumina HiSeq 2500 platform to generate 36-base single-end reads.

#### Quality control and cell selection

Prior to library construction, each capture chamber on the Fluidigm C1 chip was examined microscopically to identify wells containing exactly one captured cell, and the identifiers of these chambers were recorded. During downstream data analysis, sequencing datasets were filtered to include only libraries corresponding to these pre-annotated single-cell capture sites. For these selected cells, the read counts of three exogenous spike-in RNAs were quantified, and only cells in which the counts for all three spike-ins fell within the mean ± 2 standard deviations of their respective distributions were retained. This two-step filtering resulted in 77 HSV-1-infected cells and 84 mock-infected cells used for subsequent analyses.

#### Read alignment

Reads were first depleted of ribosomal RNA by alignment to an rRNA reference using Bowtie2 (v2.3.2). The remaining non-rRNA reads were then aligned independently to the human reference genome (hg38; -g 1 option) and to the HSV-1 reference genome (GU734771.1; -g 2 option) using TopHat2 (v2.1.1).

#### Differential expression analysis

Host differentially expressed genes (DEGs) between mock- and HSV-1-infected cells were identified with Cufflinks (v2.2.1), using only the BAM files aligned to the human genome; reads aligned to HSV-1 were excluded. Of the 6,599 genes tested, transcripts with q-values > 0.05 and –1 < log₂(fold-change) < 1 were classified as non-DEGs, yielding a total of 1,566 genes.

#### Correlation between viral load and host transcript levels

For each cell, the proportion of reads mapping to the HSV-1 genome was calculated. Spearman’s rank correlation coefficients were then computed between this viral-read percentage and the FPKM values of each non-DEG, as reported by Cufflinks.

#### Quantification of viral gene classes

Viral reads were quantified with featureCounts (Subread package) against HSV-1 coding sequences (CDS), using the parameters GTF.featureType = “CDS”, GTF.attrType = “gene”, useMetaFeatures = TRUE, and allowMultiOverlap = FALSE to exclude reads overlapping CDS from different genes. Gene-level counts were aggregated into immediate-early (IE), early (E), and late (L) classes according to HSV-1 annotations ^45^, and the proportion of each class was calculated as a percentage of the total reads assigned to the IE, E, and L genes per cell.

### Reanalysis of published scRNA-seq data from HSV-1–infected human brain organoids

A published scRNA-seq dataset of HSV-1–infected human brain organoids (GSE163952) ^23^ was reanalyzed using R with the Seurat package. The processed Seurat object (GSE163952_all_timepoints.rds.gz) provided by the original authors was used as input. Raw UMI count matrices from the RNA_exonic assay were used. For each cell, viral UMI counts were calculated by summing counts mapped to HSV-1 coding sequences. Host UMI counts were defined as total UMI counts minus viral UMI counts, and the viral fraction was calculated as viral UMI / total UMI. The analysis focused on cells harvested at 3 days post-infection. To examine the relationship between viral load and CALM gene expression while mitigating the influence of cell-to-cell variation in sequencing depth, a per-cell expression ratio was calculated for each CALM gene (CALM1, CALM2, and CALM3) as UMI_CALM / total host UMI. Cells were then divided into 100 equally spaced bins based on their viral fraction (range: 0 to 1). For each bin containing at least 30 cells, the mean viral fraction and the mean expression ratio were calculated. Spearman’s rank correlation coefficients were computed across bins to assess the association between viral load and CALM gene expression (S-Fig. 1G to I).

### Establishment of cell lines

HeLa cells were transfected with px459-CALM1 using a NEPA21 electroporator (NepaGene, Chiba, Japan). Two days after transfection, the culture medium was supplemented with 10 μg/ml puromycin overnight to select transfected cells. Puromycin was then removed, and surviving cell clones were expanded to establish CALM1-knockout (KO) cell lines. To verify genome editing, PCR products encompassing the CALM1 gRNA target site were cloned into plasmids and sequenced, revealing two distinct mutant sequences without detection of the wild-type allele (S-Fig. 2A). CALM1-KO cells were then transfected with px458-CALM2 using the same procedure. Two days post-transfection, GFP-positive cells were sorted using a FACS Melody (BD Biosciences), and resulting cell clones were expanded to establish CALM1/2-KO cell lines. Sequencing of cloned PCR products covering the CALM2 gRNA target site similarly revealed two distinct mutant sequences, with no wild-type sequence detected (S-Fig. 2B). For establishment of CALM1-KO cells stably expressing Flag-tagged calmodulin, Plat-GP cells were co-transfected with pMXs-puro-FlagCaMv2 and pMDG ^39^ using polyethylenimine (PEI), and culture supernatants were collected at 48 h post-transfection. The retrovirus-containing supernatants were added to CALM1-KO cells, followed by selection with 2 μg/ml puromycin, and resulting cell clones were designated as CALM1-KO/FlagCaM cells. Similarly, to generate HeLa cells stably expressing Flag-tagged calmodulin (HeLa/FlagCaM), Plat-GP cells were co-transfected with pMXs-puro-FlagCaMv1 and pMDG using the same method, and the resulting supernatants were used to transduce wild-type HeLa cells. After selection with 2 μg/ml puromycin, resulting cell clones were isolated and expanded. Control cell lines were also generated by transducing wild-type HeLa and CALM1-KO cells with supernatants from Plat-GP cells transfected with pMXs-puro and pMDG, followed by selection with 2 μg/ml puromycin. These lines were designated as WT/puroR and CALM1-KO/puroR cells, respectively.

### Antibodies

Commercial antibodies used in this study included mouse monoclonal antibodies to Flag (M2; Sigma), α-tubulin (DM1A; Sigma), gD (sc-21719; Santa Cruz), and ICP4 (58S; ATCC); a rabbit monoclonal antibody to calmodulin (ab45689; Abcam); and rabbit polyclonal antibodies to green fluorescent protein (GFP) (598; Medical & Biological Laboratories [MBL]) and TagRFP (AB233; Evrogen). Rabbit polyclonal antibodies to ICP47, Us11, and UL12 were generated in-house in our previous studies ^22,46,47^.

### Immunoblotting

Cell lysates in SDS sample buffer (62.5 mM Tris–HCl pH 6.5, 20% glycerol, 2% SDS and 5% 2-mercaptoethanol) were sonicated, heated (100 °C), subjected to electrophoresis in denaturing gels and transferred to PVDF membranes (Merck). The membranes were blocked with 5% skim milk in T-PBS (PBS containing 0.05% Tween 20) for at least 30 min, followed by incubation with the indicated antibodies for at least 2 h at room temperature or 4 °C. These membranes were then incubated with secondary antibodies conjugated with peroxidase (Cytiva) for at least 1 h at room temperature or 4 °C and visualized using ECL (Cytiva) with an ImageQuant LAS 4000 (GE Healthcare) or a ChemiDoc MP (Bio-Rad) system. The protein present in immunoblot bands was quantified using the ImageQuant LAS 4000 system with the ImageQuant TL7.0 analysis software (GE Healthcare) or ChemiDoc MP with the Image Lab 6.01 software (Bio-Rad) according to the manufacturer’s instructions and normalized to that of α-tubulin proteins.

### Flow cytometry

HeLa cells infected with the indicated viruses under the described experimental conditions were washed once with phosphate-buffered saline (PBS) and detached using 0.25% trypsin–EDTA solution (Fujifilm). The cells were then resuspended in PBS containing 2% fetal calf serum (FCS) and analyzed on either a BD FACSMelody (Becton Dickinson) or a CytoFLEX S (Beckman Coulter) flow cytometer. The acquired data were processed and analyzed using FlowJo software version 10.10.0 (Becton Dickinson).

### Definition of gating for flow cytometry analysis

Gates to distinguish positive and negative populations for each fluorescent protein were established by simultaneously analyzing cells infected with wild-type HSV-1(F), which served as a negative control. The Venus^high^ cell population was defined by first sorting the f4–f6 subpopulations—previously identified in our earlier study as the principal producers of progeny virus—under the flow cytometry settings used in that study ^22^. These sorted populations were then immediately reanalyzed on the same day under the flow cytometry conditions employed in the present study, and the Venus^high^ gate was drawn to include events whose Venus fluorescence intensity was equal to or greater than that of the f4 subpopulation.

For experiments measuring viral titers from sorted cells, additional gates were applied to subdivide cells based on Venus fluorescence intensity. Specifically, the low-1 and low-2 subpopulations were defined as cells with Venus fluorescence intensities lower than those of the Venus^high^ gate, subdivided into two bins of the fluorescence intensity histogram. The Venus^high^ population was further split into high-1 and high-2: high-1 corresponded to the f4 subpopulation in our previous study, whereas high-2 encompassed the range corresponding to the f5–f6 subpopulations. Thus, a total of four subpopulations (low-1, low-2, high-1, and high-2) were defined for sorting and subsequent viral titer determination.

### Quantitative reverse transcription PCR (qRT-PCR)

Total RNA was isolated from infected cells, and cDNA was synthesized as described previously ^48^. The amounts of cDNA for specific genes were quantified using the Universal Probe Library (Roche) together with the TaqMan Master Mix (Roche) on a LightCycler 96 system (Roche), according to the manufacturer’s instructions. Gene-specific primers and universal probes were designed using the ProbeFinder software (Roche). The primer sequences and probes used in this study are listed in Table S3. The levels of the indicated mRNAs were calculated either as 2−^ΔCt^, where ΔCt represents the difference between the Ct values of the target gene and 18S rRNA, or as relative copy numbers determined by absolute quantification. For absolute quantification, standard curves were generated using serial dilutions of plasmids (pFLAG-UL29, pFLAG-ICP0, pFLAG-UL54, pFLAG-Us2, pFLAG-UL49, pFLAG-UL50, or pFLAG-18S), and the copy numbers of cDNA derived from each mRNA were determined based on these curves. The copy number of each target gene was then normalized to that of 18S rRNA to obtain relative expression levels.

### Determination of fluorescent protein expression onset by time-lapse imaging

A total of 2.5 × 10^5^ indicated cells were seeded into collagen-coated 24-well glass-bottom dishes. Prior to seeding, 10% of the cells were labeled with CellTrace Far Red (Thermo) according to the manufacturer’s instructions and then mixed with 90% unlabeled cells. Six hours after seeding, cells were infected with the indicated viruses at an MOI of 5 by adsorption for 1 h at 37°C in 250 µl of 199 medium supplemented with 1% FCS. The inoculum was then replaced with 500 µl of clear 199 medium (Thermo) containing 1% FCS.

Time-lapse fluorescence images were acquired every 15 min from 1.5 hpi up to 24 hpi using a BZ-X810 microscope (Keyence) with a 10x objective lens (Plan-Apochromat, NA 0.45), equipped with a stage top incubator (TOKAI HIT). The Cy5 filter set (excitation 620/60 nm, emission 700/75) was used to detect Far Red, the mVenus filter set (excitation 500/20 nm, emission 535/30) for Venus, and the TagRFP filter set (excitation 546/22 nm, emission 590/33) for TagRFP.

All image analyses were performed using ImageJ (version 1.54). To avoid signal saturation during subsequent spectral unmixing, the intensities of all channels were first reduced by half. Background subtraction was then applied to all images using the rolling ball method with a radius of 50. Spectral unmixing was performed using the Spectral Unmixing plugin (https://imagej.net/ij/plugins/spectral-unmixing.html) to separate Venus and TagRFP signals. The unmixing matrix was calculated from reference images of control cells displaying fluorescence solely from TagRFP or Venus.

Cell segmentation was conducted using the Cellpose detector in TrackMate with the cyto3 model ^24,49^, applied to the Far Red channel with an estimated cell diameter parameter of 27.2 µm. Tracking was carried out using the LAP tracker with a maximum linking distance of 14 µm and an area feature penalty of 1.0, without enabling gap closing, splitting, or merging.

Fluorescence onset analysis was performed on cells that were continuously tracked from the beginning to the end of time-lapse imaging. Tracking data exported from TrackMate were analyzed in R (version 4.4.1) using custom scripts. For each tracked cell, mean fluorescence intensities of TagRFP and Venus were extracted at each time point. The baseline fluorescence for each channel was calculated as the mean intensity over the first five time points. A fluorescence onset was defined as the first time point at which the mean intensity exceeded the baseline plus five times the standard deviation of the baseline, sustained for at least five consecutive time points. The onsets for TagRFP and Venus were then recorded as the corresponding time in minutes, and the time lag between Venus and TagRFP onset was calculated.

### Direct Calmodulin–ICP4 Binding Assay

293FT cells were transfected with pFLAG-ICP4, pFLAG-ICP4ΔIQ, or pFLAG-EGFP plasmids using polyethyleneimine (PEI). Two days post-transfection, cells were harvested and lysed on ice for 20 min in NP40 buffer (120 mM NaCl, 50 mM Tris-HCl [pH 8.0], 0.5% NP-40, 50 mM NaF) supplemented with protease inhibitor cocktail (Nacalai Tesque). The lysates were centrifuged to remove cell debris, and the resulting supernatants were incubated with anti-FLAG affinity gel (Sigma) for 2 h at 4°C with rotation. After centrifugation, the supernatants were discarded, and the affinity gels were washed four times with NP40 buffer followed by a single wash with high-salt NP40 buffer (300 mM NaCl, 50 mM Tris-HCl [pH 8.0], 0.5% NP-40, 50 mM NaF). The gels were then resuspended in FLAG elution buffer (50 mM Tris-HCl [pH 8.0], 170 mM NaCl, 0.5 mg/ml FLAG peptide) and rotated for 2 h at 4°C. After centrifugation, the eluates were collected and dialyzed against elution buffer lacking FLAG peptide to remove excess FLAG peptide. The resulting samples were used as purified FLAG-tagged proteins for subsequent binding assays.

For the binding assay, 50 μl (50% slurry) of either Calmodulin Sepharose 4A beads (GE Healthcare) or Protein A Sepharose 4A beads (GE Healthcare) was added to 500 μl of binding buffer (50 mM Tris-HCl [pH 8.0], 150 mM NaCl, 1 mM CaCl₂, 0.05% NP-40) supplemented with protease inhibitor cocktail (Nacalai Tesque). Purified FLAG-tagged proteins (500 ng each) were then added to the bead-containing buffer and incubated at 30°C for 1 h with rotation. After incubation, the beads were washed three times with NP40 buffer, resuspended in sample buffer, and subjected to SDS-PAGE.

### Immunoprecipitation

HeLa/puroR or HeLa/Flag-CaM cells were infected with wild-type HSV-1(F) at a multiplicity of infection (MOI) of 5. At 9 h post-infection, cells were harvested and lysed in NP-40 buffer (120 mM NaCl, 50 mM Tris-HCl [pH 8.0], 0.5% NP-40, and 50 mM NaF) supplemented with a protease inhibitor cocktail and a phosphatase inhibitor cocktail (both from Nacalai Tesque). Lysates were clarified by centrifugation and passed through a 0.2-µm filter (Sartorius), followed by incubation with Dynabeads Protein G (Thermo Fisher Scientific) pre-conjugated with anti-Flag M2 antibody. The mixtures were rotated overnight at 4 °C. After extensive washing with NP-40 buffer, bound proteins were eluted in SDS sample buffer and subjected to SDS–PAGE.

### ChIP-qPCR

ChIP-qPCR was performed as described previously ^50^. Briefly, HeLa cells were infected with either YK416 (ICP4ΔIQ) or YK417 (ICP4ΔIQ-repair) at an MOI of 5, or HeLa and HeLa/CALM1-KO cells were infected with wild-type HSV-1(F) at an MOI of 5. At 3.5 h post-infection, cells were fixed, treated with lysis buffer as described previously ^50^, and sonicated 15 times with 30 s pulses using a sonicator (UR-21P; TOMY), followed by the addition of Triton X-100 to a final concentration of 1%. After centrifugation, supernatants were precleared by rotation with Dynabeads protein G (Invitrogen), and immunoprecipitated with anti-ICP4 antibody (58S; ATCC) or control IgG (MOPC-173; BioLegend) bound to Dynabeads protein G at 4°C overnight. The beads were washed eight times with ChIP-RIPA buffer [50 mM HEPES-KOH (pH 7.5), 500 mM LiCl, 1 mM EDTA, 1% NP-40, 0.7% sodium deoxycholate] and once with TNE buffer [50 mM Tris-HCl (pH 8.0), 10 mM EDTA, 50 mM NaCl]. DNA-protein complexes were eluted, reverse cross-linked, and treated with RNase A and Proteinase K as described previously ^50^. DNA was purified using a NucleoSpin gel and PCR clean-up kit (Machery-Nagel) according to the manufacturer’s instructions. qPCR was carried out with SYBR Green on a LightCycler 96 system (Roche) using primer sets listed in Table S4. The ΔCt was calculated by subtracting the Ct values of ChIP DNA from the Ct values of input DNA and the relative amount of ChIP DNA to input DNA (2−ΔCt) for each sample was normalized to the sum of all values in one experiment (relative enrichment).

### Determination of viral titer in the subpopulations separated by cell sorting

Determination of viral titers from sorted subpopulations was performed as described previously ^22^ with the following modifications. The indicated cell lines were infected with the indicated viruses at an MOI of 5. At the indicated times post-infection, cells were washed with phosphate-buffered saline (PBS) and detached with 0.25% trypsin–EDTA solution (Fujifilm). The cells were then resuspended in PBS containing 2% fetal calf serum (FCS) (washing buffer), passed through a 35-µm cell strainer (#352235, Corning), and subjected to fluorescence-activated cell sorting (FACS) on a BD FACS Melody (Becton Dickinson). A total of 3 × 10^3^ cells from each of the four subpopulations (low-1, low-2, high-1, and high-2), as well as from the FSC-singlet population, were collected into medium 199 supplemented with 1% FCS. The sorted cells underwent a single freeze–thaw cycle followed by sonication, and viral titers were determined by plaque assay on Vero cells. The average viral yield per cell was calculated by dividing the total titer by the number of sorted cells. Data are expressed as PFU per 10^4^ cells.

### Calculation of the virtual titer

The virtual titer (VT) is defined as the sum of subpopulation-specific titers (PFU_subi_), representing the estimated viral titer of the entire cell population reconstructed from subpopulation data. Virtual titers were calculated essentially as described previously^22^, with subpopulations defined in the present study (low-1, low-2, high-1, and high-2). The PFU for each subpopulation (PFU_subi,_ where subi = low-1, low-2, high-1, and high-2) was calculated using the following equation:

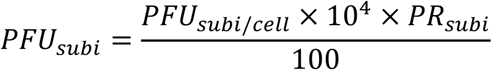

where PFU_subi/cell_ is the average viral titer per single cell belonging to the subi subpopulation, and PR_subi_ is the proportion (%) of cells in that subpopulation within the entire population (FSC singlet).

The virtual titer (VT), representing the sum of all PFU_subi_ values, was obtained as follows:

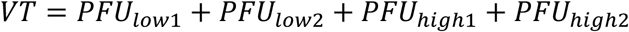

The percentage contribution of each subpopulation to the total viral titer (% of PFU_subi_) was calculated as:

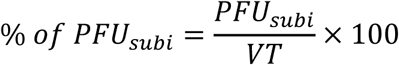

PFU_subi_ values for each subpopulation were calculated as described above, using PFU_subi/cell_ and PR_subi_ obtained from the same sample. In addition, a cross-combination analysis was performed in which PFU_subi/cell_ values obtained from one sample were combined with PR_subi_ values obtained from another sample.

### Animal studies

Three-week-old female ICR mice were purchased from Charles River Laboratories and used for the experiments. Mice were inoculated intracranially with 5 × 10^2^ PFU of wild-type HSV-1(F) mixed with the indicated amount of acyclovir (ACV; Fujifilm) or W-7 (Sigma-Aldrich), and survival was monitored daily for 14 days. All animal experiments were carried out in accordance with the Guidelines for Proper Conduct of Animal Experiments, Science Council of Japan. The protocol was approved by the Institutional Animal Care and Use Committee (IACUC) of the Institute of Medical Science, the University of Tokyo (protocol approval numbers: PM27-113, PM27-73, PA17-67, PH2-12, A21-55 and A22-27).

### Statistical analysis

Statistical analysis was performed as follows. Comparisons between two groups were conducted using Welch’s t-test or the Mann–Whitney U-test. For qPCR data normalized to a specific sample set as 1, a one-sample t-test was used, and p-values were adjusted by the Bonferroni correction when multiple pairwise comparisons were performed. Viral titers were log10-transformed prior to statistical testing. For comparisons among three or more groups, one-way ANOVA followed by Tukey’s multiple-comparison test or the Kruskal–Wallis test followed by Dunn’s multiple-comparison test was applied. Survival curves were compared using the log-rank test, with Holm’s correction applied when multiple pairwise comparisons were performed. For comparison of onset detection rates, Fisher’s exact test was used with Bonferroni correction. In fluorescence onset analyses, cells in which onset was not detected during the imaging period were assigned an imputed value (maximum observed onset + 60 min) and included in statistical testing. Statistical significance was defined as p < 0.05. The statistical tests used for each figure are indicated in the corresponding figure legends.

## ACKNOWLEDGEMENTS

We thank Kiyomi Imamura, Shota Takashima, Tomohiko Takeda, Risa Fujinaga, Risa Abe, Tohru Ikegami and Yui Muto for their excellent technical assistance.

## Funding

This study was supported by Grants for Scientific Research and Grant-in-Aid for Scientific Research (S) (20H05692) from the Japan Society for the Promotion of Science (JSPS), a PRESTO grant (JPMJPR22R5) from the Japan Science and Technology Agency (JST), grants (JP20wm0125002, JP22gm1610008, JP223fa627001, JP23wm0225031, and JP23wm0225035) from the Japan Agency for Medical Research and Development (AMED), grants from the International Joint Research Project of the Institute of Medical Science, the University of Tokyo, and grants from the Takeda Science Foundation, the Uehara Memorial Foundation, Shionogi Infectious Disease Research Promotion Foundation, and the Morinomiyako Medical Research Foundation.

## Author contributions

Conceptualization: Y.M. and Y.Ka.; Methodology: Y.M. and Y.S.; Investigation: Y.M., M.S., and Y.Ku.; Resources: Y.M., A.K., and Y.S.; Visualization: Y.M.; Funding acquisition: Y.M., A.K., Y.S., and Y.Ka.; Writing – original draft: Y.M. and Y.Ka.; Writing – review & editing: All authors; Project administration: Y.S. and Y.Ka.; Supervision: Y.S. and Y.Ka.

## Competing interests

Y.M. and Y.Ka. are inventors on a patent application related to the analysis method used in this study (Japanese Patent Application No. 2025-104075, filed June 19, 2025). The other authors declare no competing interests.

## Data and code availability

The single-cell RNA-sequencing data generated in this study have been deposited at the DNA Data Bank of Japan (DDBJ) and are publicly available. Raw sequencing reads and processed gene-expression matrices are available at the DDBJ Sequence Read Archive (DRA) and the Genomic Expression Archive (GEA) under accession numbers DRR1056414–DRR1056574 and E-GEAD-1262, respectively. Custom R scripts used for fluorescence onset analysis are available from the corresponding author upon request. Any additional information required to reanalyze the data reported in this paper is available from the corresponding author upon request.

## Declaration of generative AI and AI-assisted technologies in the manuscript preparation process

During the preparation of this work, the authors used Claude (Anthropic) and ChatGPT (OpenAI) in order to improve the language and readability of the manuscript. After using these tools, the authors reviewed and edited the content as needed and take full responsibility for the content of the publication.

## Notes

### Summary of Updates

Abstract, Introduction, Results, and Discussion updated to clarify the wording.

