## Supplementary Figures 1-10 for "Calmodulin controls the tempo of HSV-1 gene-expression cascade to reshape infection heterogeneity"

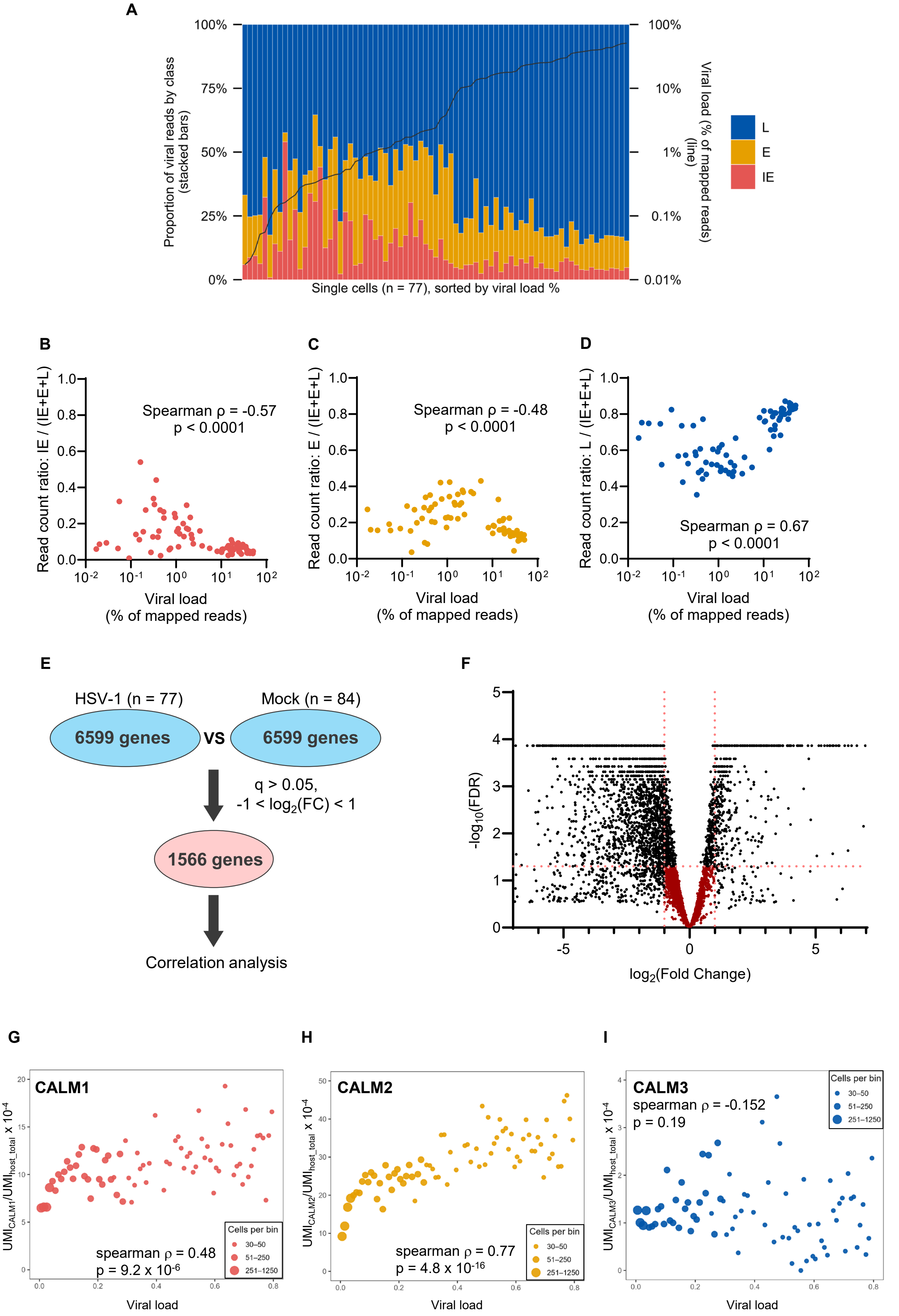


*(legend on next page )*

**S-Fig. 1 Y. Maruzuru et al.**

**S-Fig. 1. Correlation of CALM gene expression with viral load and gene class composition in single cells.**
(A) Stacked bar graph showing the proportion of IE, E, and L reads relative to their sum (IE + E + L) for each of the 77 HSV-1–infected single cells (left y-axis), sorted by viral load. The overlaid line indicates the viral load (% of total mapped reads) for each cell (right y-axis). Cells were infected at an MOI of 5 and harvested at 13 h post-infection for library preparation.
(B to D) Scatter plots showing the relationship between the viral load and the proportion of (B) IE, (C) E, and (D) L reads relative to their sum (IE + E + L) in each single cell. Spearman's rank correlation coefficients (ρ) and p values are indicated. (E) Schematic overview of the filtering strategy for correlation analysis. Among 6,599 host genes tested for differential expression, 1,566 genes whose expression was not significantly altered between HSV-1-infected and mock-infected cells ($q > 0.05$, $-1 < \log_2 FC < 1$) were selected and used for correlation analysis with the fraction of viral reads. (F) Volcano plot of host genes (out of 6,599 tested) showing $\log_2$(Fold Change) versus $-\log_{10}$(FDR) between HSV-1-infected ($n = 77$) and mock-infected ($n = 84$) cells. Genes within the dashed lines ($q > 0.05$, $-1 < \log_2 FC < 1$) represent the 1,566 genes used for correlation analysis as shown in Fig. 1A.
(G to I) Reanalysis of a published scRNA-seq dataset of HSV-1–infected human brain organoids [23]. Scatter plots show the correlation between average viral load per bin and average UMI ratio (UMI of each CALM gene / total host UMI $\times 10^{-4}$) per bin for (G) CALM1, (H) CALM2, and (I) CALM3. Cells were grouped into bins based on their viral load; dot size indicates the number of cells per bin. Spearman's rank correlation coefficients (ρ) and p values are indicated.

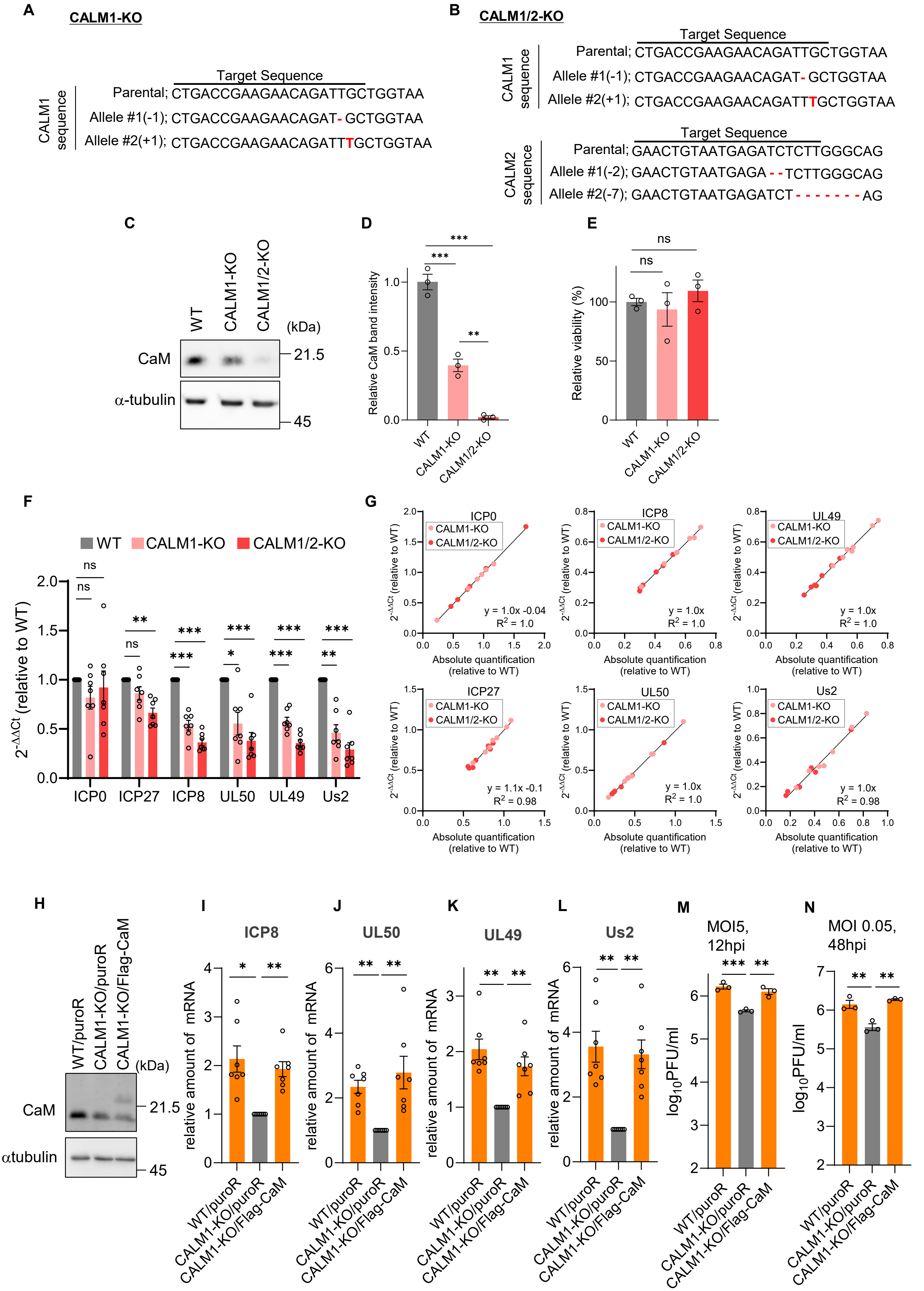


*(legend on next page )*

**S-Fig. 2 Y. Maruzuru et al.**

**S-Fig. 2. Generation of CaM-deficient cells and characterization of rescue cell lines.**
(A and B) Genomic DNA sequences of the CRISPR-Cas9 target sites in CALM1-KO cells (A) and CALM1/2-KO cells (B). The parental (wild-type) sequence and the identified mutant alleles (insertions/deletions, indicated by dashes or red letters) are shown for each gene.
(C) Immunoblot analysis of calmodulin (CaM) in HeLa wild-type (WT), CALM1-knockout (CALM1-KO), and CALM1/2-knockout (CALM1/2-KO) cells. α-tubulin served as a loading control.
(D) Quantification of CaM band intensities normalized to α-tubulin.
(E) Cell viability of WT, CALM1-KO, and CALM1/2-KO cells assessed by a WST-8 assay.
(F) Relative mRNA levels calculated using the ΔΔCt method from the same raw data shown in Fig. 2A. HeLa WT, CALM1-KO, and CALM1/2-KO cells were infected with wild-type HSV-1(F) at an MOI of 5. At 6 h post-infection, total RNA was extracted, and the resulting data were analyzed. Data are shown relative to WT levels (set to 1).
(G) Correlation plots comparing the relative mRNA levels obtained by the absolute quantification method (Fig. 2A, x-axis) and the ΔΔCt method (S-Fig. 2F, y-axis) for each viral gene. Linear regression lines and $R^2$ values are shown.
(H) Immunoblot analysis of CaM in control (WT/puroR, CALM1-KO/puroR) and Flag-CaM complemented (CALM1-KO/Flag-CaM) cell lines. α-tubulin served as a loading control.
(I to L) WT/puroR, CALM1-KO/puroR, CALM1-KO/Flag-CaM cells were infected with wild-type HSV-1(F) at an MOI of 5 for 6 h. Relative mRNA levels of E genes (I: ICP8, J: UL50) and L genes (K: UL49, L: Us2) were quantified by qRT-PCR using the ΔΔCt method.
(M and N) WT/puroR, CALM1-KO/puroR, CALM1-KO/Flag-CaM cells were infected with wild-type HSV-1(F) at an MOI of 5 for 12 h (M) or at an MOI of 0.05 for 48 h (N). Progeny virus titers were determined by plaque assay. Data are representative of three independent experiments (C and H). Data are presented as mean ± SE of seven (F, I to L), or three (D, E, M, and N) biological replicates. Statistical analyses were performed using a one-sample t-test with Bonferroni correction (F, I to L) or one-way ANOVA followed by Tukey's multiple-comparison test (D, E, M, and N). *, $p < 0.05$; **, $p < 0.01$; ***, $p < 0.001$; ****, $p < 0.0001$; ns, not significant.

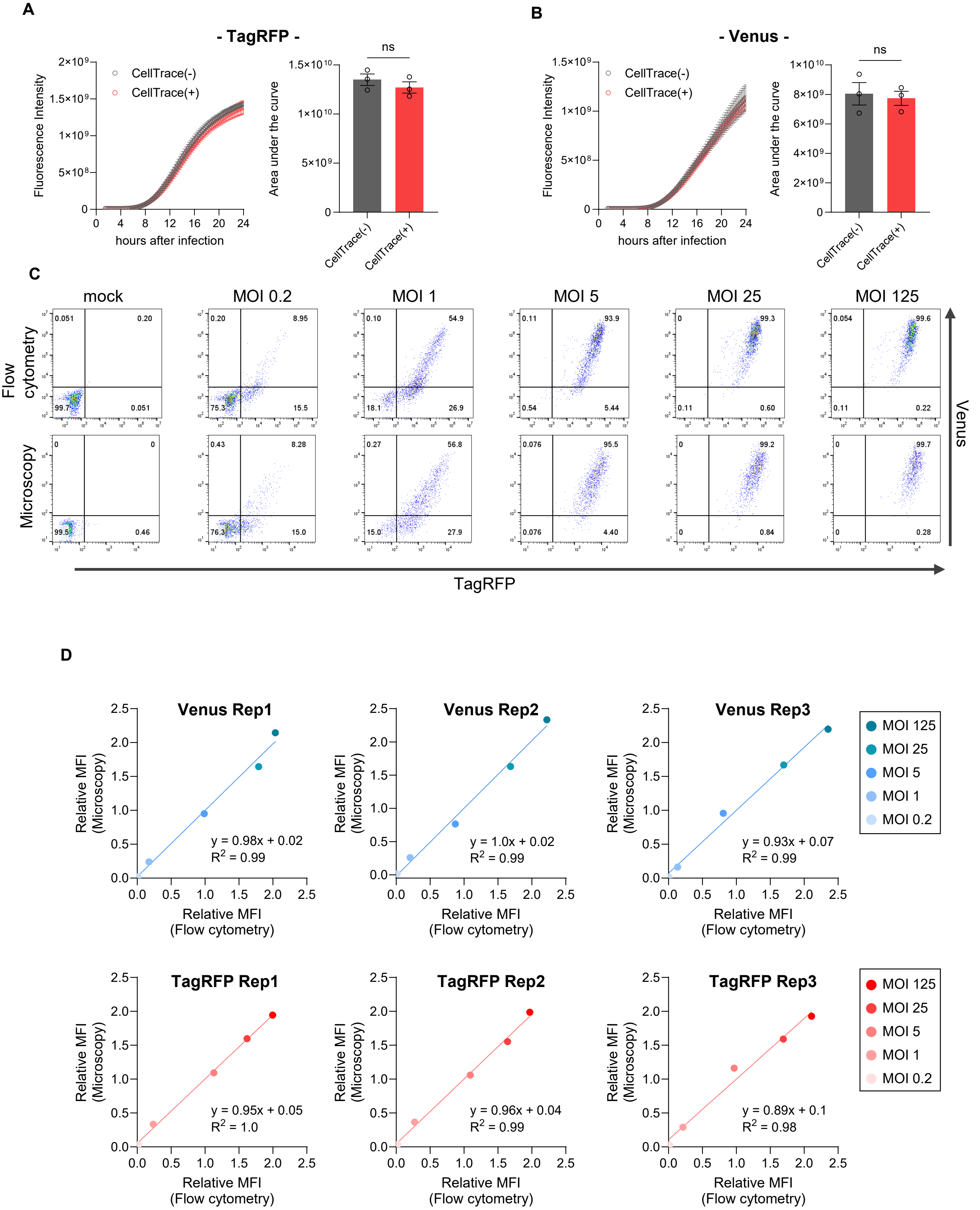


*(legend on next page )*

**S-Fig. 3 Y. Maruzuru et al.**

**S-Fig. 3. Effect of CellTrace labeling on viral fluorescence accumulation and comparison of single-cell fluorescence quantification with flow cytometry.**

(A and B) HeLa cells were labeled with CellTrace Far Red (CellTrace$^{+}$) or left unlabeled (CellTrace$^{-}$), seeded in separate wells, and infected with rICP47/vUs11 (MOI of 5). Time-lapse imaging was performed to simultaneously monitor TagRFP (IE) and Venus (L) expression in the same fields of view. (A) TagRFP (IE). (B) Venus (L). Left panels show total fluorescence intensity per field over time; right panels show the area under the curve (AUC) from 1.5 to 24 h post-infection (hpi). Data represent mean ± SE from three independent experiments. For each experiment, the total fluorescence intensity per field was averaged across three imaging fields and used as a representative value. Statistical analysis (right panels) was performed using Welch's t-test. ns, not significant.

(C) HeLa cells (a 1:9 mixture of CellTrace Far Red–stained and unstained cells) were infected with rICP47/vUs11 at the indicated MOIs (0.2, 1, 5, 25, 125) or mock-infected. At 16 h post-infection, the same wells were first imaged by microscopy and then analyzed by flow cytometry. Representative TagRFP vs. Venus fluorescence plots comparing both methods are shown. The top panels show data from flow cytometry; the bottom panels show corresponding microscopy data, where the fluorescence intensity of each segmented CellTrace$^{+}$ cell was plotted (as in Fig. 3A).

(D) Correlation analysis comparing microscopy- and flow cytometry–based quantification from the experiment in (C). The relative mean fluorescence intensity (MFI) measured by microscopy (y-axis) was plotted against that measured by flow cytometry (x-axis). The top row and bottom row show correlations for Venus (L) and TagRFP (IE), respectively. Each point represents the average fluorescence intensity of cells from the same sample, collected across infections at different MOIs (color-coded). Results from three independent experiments (Rep1, Rep2, and Rep3) are shown, each with the corresponding linear regression line and $R^2$ value.

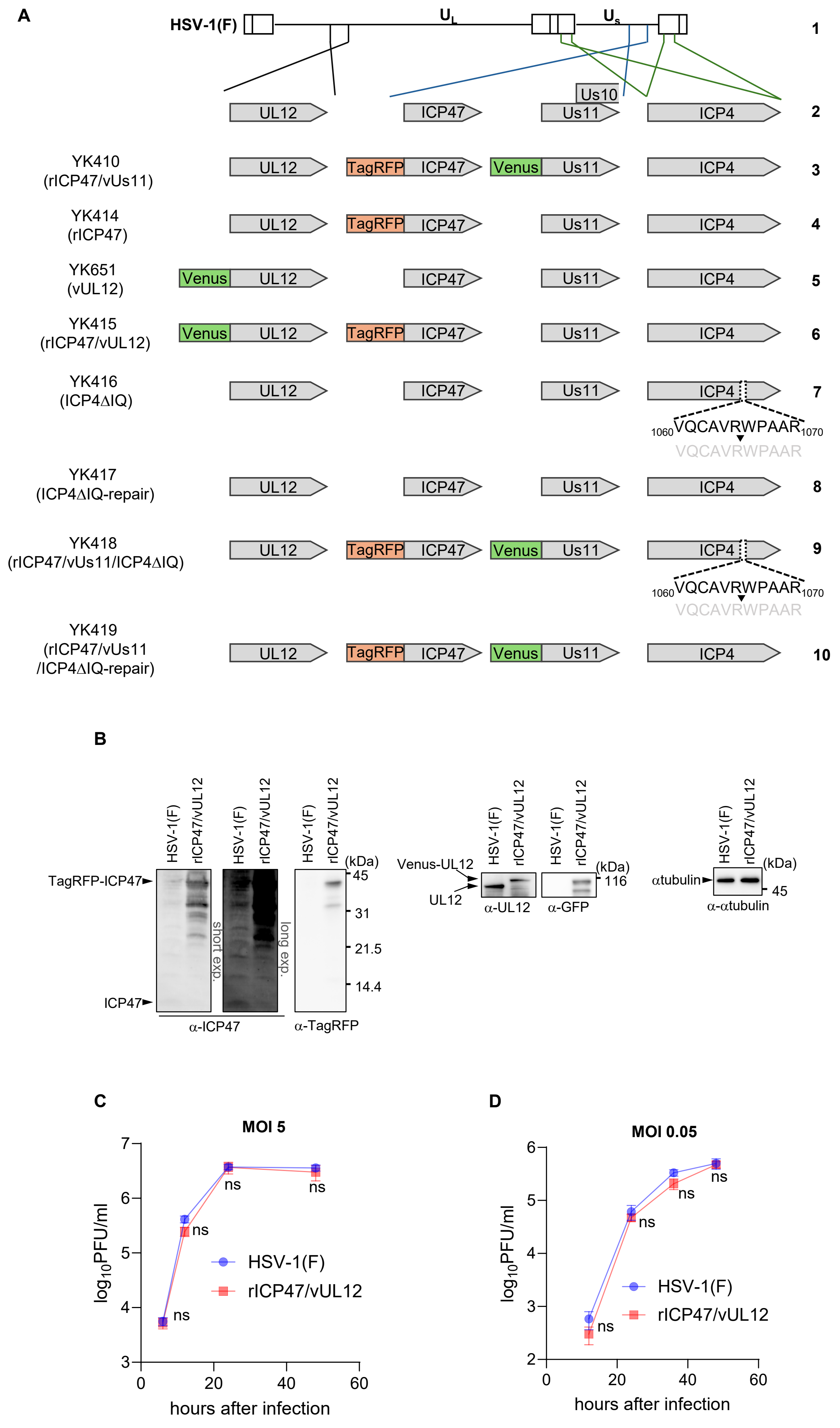


*(legend on next page )*

**S-Fig. 4 Y. Maruzuru et al.**

**S-Fig. 4. Generation and characterization of recombinant HSV-1 reporter viruses.**
(A) Schematic diagram of the genomic structures of wild-type HSV-1(F) and recombinant viruses used in this study. Line 1, wild-type HSV-1(F) genome; line 2, domains of the genes encoding UL12, ICP47, Us11, Us10, and ICP4; lines 3–10, recombinant viruses used in this study. Fluorescent protein tags (TagRFP or Venus) are indicated.
(B) HeLa cells were infected with wild-type HSV-1(F) or rICP47/vUL12 at an MOI of 5. Cell lysates were harvested at 24 h post-infection and analyzed by immunoblotting with the indicated antibodies.
(C and D) Viral growth kinetics of reporter viruses. HeLa cells were infected with wild-type HSV-1(F) or rICP47/vUL12 at an MOI of 5 (C) or 0.05 (D). At the indicated times post-infection, infected cells and culture supernatants were harvested together, and progeny virus titers were determined by plaque assay on Vero cells.
Data represent the mean ± SE of three biological replicates. Statistical analysis was performed using Welch's t-test. ns, not significant.

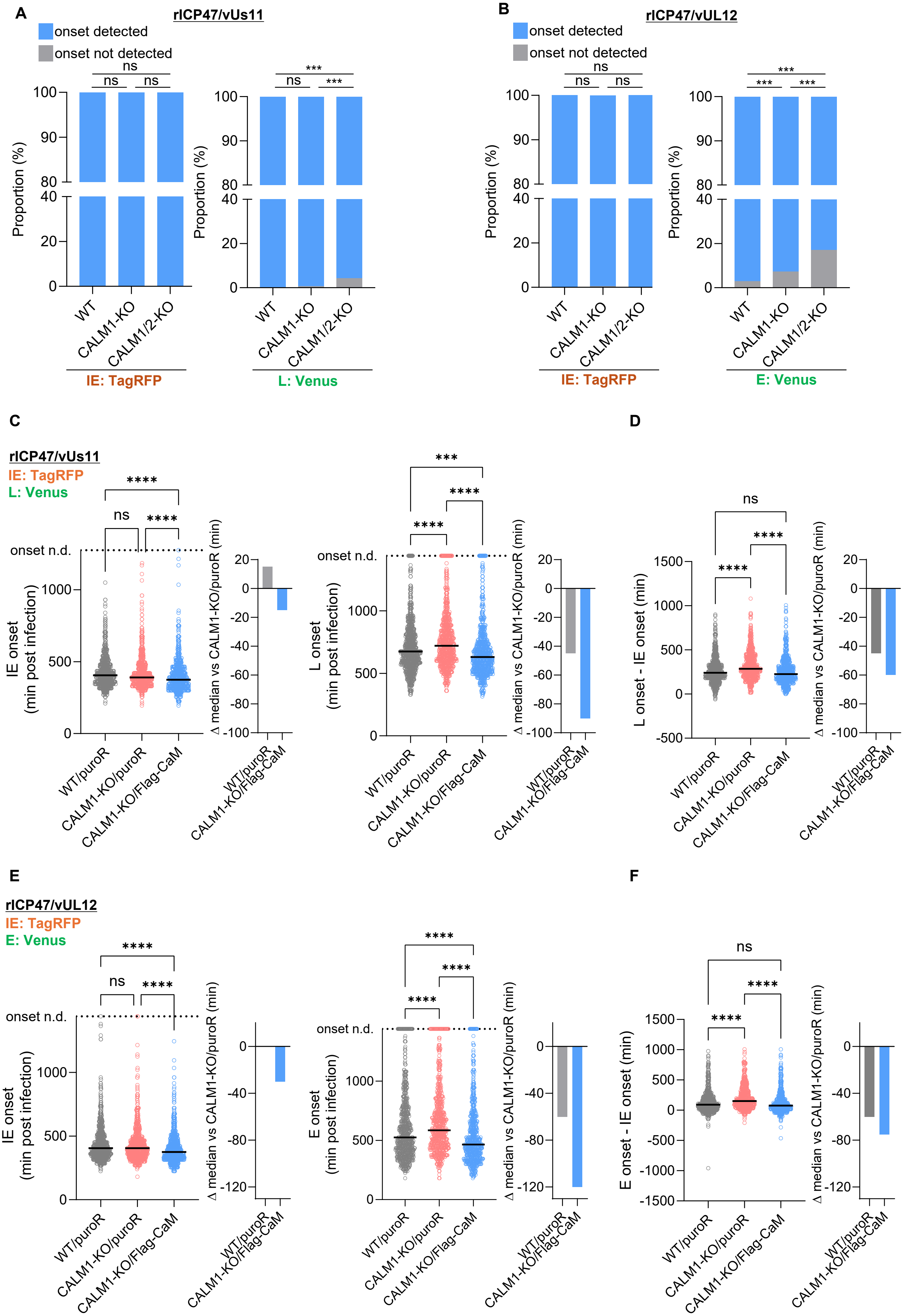


*(legend on next page )*

**S-Fig. 5 Y. Maruzuru et al.**

**S-Fig. 5. CaM deficiency delays the IE-to-E/L transition, which is rescued by CaM complementation.**

(A and B) HeLa WT, CALM1-KO, and CALM1/2-KO cells were infected with rICP47/vUs11 (A) or rICP47/vUL12 (B) at an MOI of 5 and imaged every 15 min from 1.5 to 24 h post-infection. The stacked bar graphs indicate the percentages of tracked cells in which the onset of TagRFP and Venus expression was either detected or not detected during the observation period. These data are derived from the single-cell tracking analysis shown in Fig. 3B–E. Statistical analyses were performed using Fisher's exact test with Bonferroni correction; ***, $p < 0.001$; ns, not significant.

(C and D) Control (WT/puroR, CALM1-KO/puroR) and Flag-CaM-complemented CALM1-KO (CALM1-KO/Flag-CaM) cells were infected with rICP47/vUs11 at an MOI of 5 and imaged every 15 min from 1.5 to 24 h post-infection. (C) Dot plots showing IE onset (TagRFP, left) and L onset (Venus, right) times (min post-infection) for individual cells. Cells in which onset was not detected are plotted above the dashed line (onset n.d.) with an imputed value (maximum observed onset + 60 min) and were included in statistical analyses. (D) Time interval between IE and L onset (L onset − IE onset) in cells where both onsets were defined.

(E and F) Control (WT/puroR, CALM1-KO/puroR) and Flag-CaM-complemented CALM1-KO (CALM1-KO/Flag-CaM) cells were infected with rICP47/vUL12 at an MOI of 5 and imaged every 15 min from 1.5 to 24 h post-infection. (E) Dot plots showing IE onset (TagRFP, left) and E onset (Venus, right) times (min post-infection) for individual cells. Cells in which onset was not detected are plotted above the dashed line (onset n.d.) with an imputed value (maximum observed onset + 60 min) and were included in statistical analyses. (F) Time interval between IE and E onset (E onset − IE onset) in cells where both onsets were defined.

In (C to F), each dot represents one cell and bars indicate medians. Adjacent bar graphs indicate the differences in median values relative to CALM1-KO/puroR. The number of analyzed cells was as follows: (C) WT/puroR, n = 783; CALM1-KO/puroR, n = 718; CALM1-KO/Flag-CaM, n = 664; (D) WT/puroR, n = 767; CALM1-KO/puroR, n = 686; CALM1-KO/Flag-CaM, n = 644; (E) WT/puroR, n = 694; CALM1-KO/puroR, n = 644; CALM1-KO/Flag-CaM, n = 626; (F) WT/puroR, n = 655; CALM1-KO/puroR, n = 604; CALM1-KO/Flag-CaM, n = 605. Data represent pooled measurements from two independent experiments. Statistical analyses were performed using the Kruskal–Wallis test followed by Dunn's multiple-comparison test; ****, $p < 0.0001$; ***, $p < 0.001$; ns, not significant.

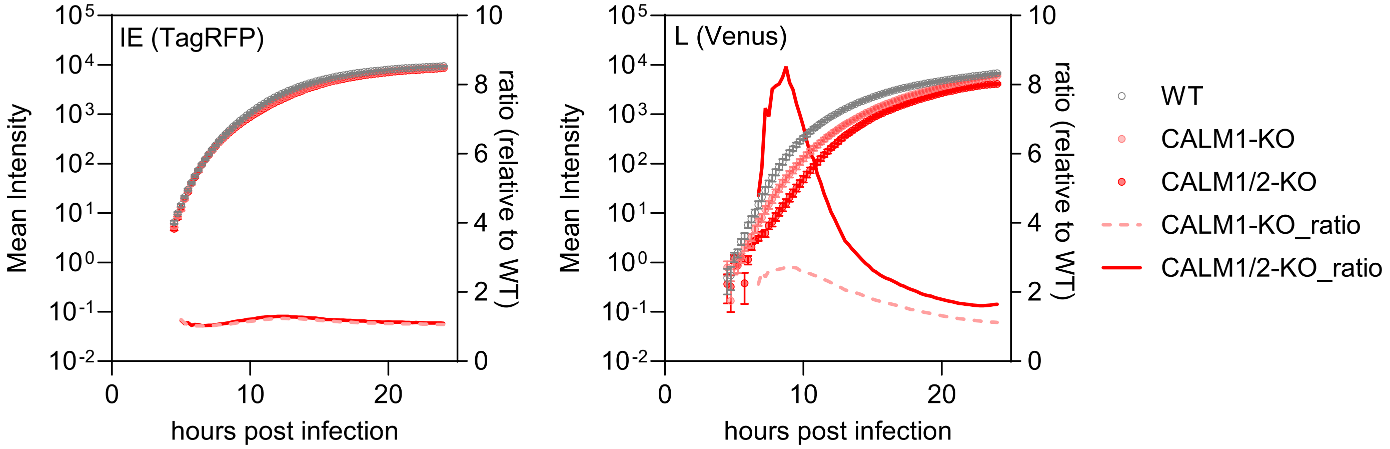


**S-Fig. 6. Delay in L protein accumulation in CaM-deficient cells.**

Mean fluorescence intensity from Fig. 4B is replotted on a logarithmic y-axis (left axis) for IE (TagRFP, left) and L (Venus, right). The ratio of mean fluorescence intensity relative to WT (CALM1-KO/WT and CALM1/2-KO/WT) is also shown (right axis). Data are presented as mean ± SE.

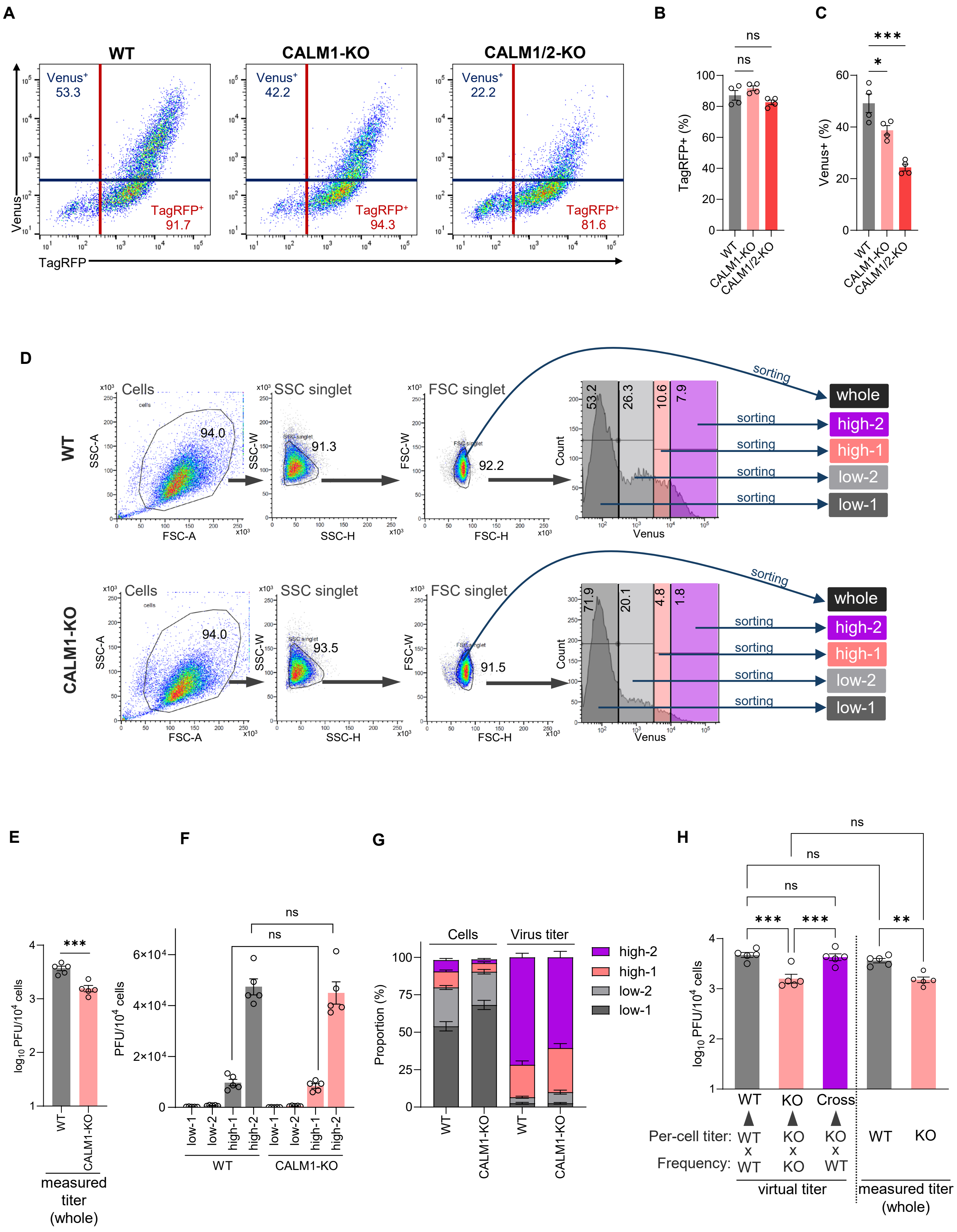


*(legend on next page )*

**S-Fig. 7 Y. Maruzuru et al.**

**S-Fig. 7. Effect of CALM1-KO on viral gene expression heterogeneity and progeny virus production.**
(A to C) HeLa WT, CALM1-KO, and CALM1/2-KO cells were infected with rICP47/vUs11 at an MOI of 5. At 12 h post-infection, cells were analyzed by flow cytometry. (A) Representative flow cytometry plots showing TagRFP (IE) and Venus (L) fluorescence. Gates for TagRFP+ and Venus+ populations are shown, with the percentage of cells within these gates indicated. (B) Quantification of the percentage of TagRFP+ (IE) cells from (A). (C) Quantification of the percentage of Venus+ (L) cells from (A).
(D to H) HeLa WT and CALM1-KO cells were infected as in (A) and subjected to fluorescence-activated cell sorting (FACS) at 12 h post-infection.
(D) Gating strategy for cell sorting. Single cells (SSC- and FSC singlet) were sorted into the whole population or fractionated into four subpopulations (low-1, low-2, high-1, high-2) based on Venus (L) fluorescence intensity.
(E) Viral titers (PFU per $10^4$ cells) from the sorted "whole" singlet population.
(F) Viral titers of each of the four sorted subpopulations in (D) (i.e., the per-cell titers used in H).
(G) Stacked bar graphs showing, for WT and CALM1-KO cells, the subpopulation composition ("Cells") and each subpopulation's contribution to the total viral yield ("Virus titer"), both expressed as percentages.
(H) Virtual titer analysis. For each Venus-defined subpopulation, the product of its frequency among infected cells ("Frequency") and its per-cell titer ("Per-cell titer") was calculated, and these products were summed across subpopulations to obtain the virtual titer. The genotype source of each parameter (WT or CALM1-KO) is indicated below each bar: the "WT" and "KO" bars use the Frequency and Per-cell titer from the same genotype, whereas the "Cross" bar combines the WT Frequency with the CALM1-KO Per-cell titer. Measured whole-population titers (as in E) are shown on the right for comparison.
Each value represents the mean ± SE of four (B and C) or five (E to H) biological replicates. Statistical analyses were performed using one-way ANOVA followed by Tukey's multiple-comparison test (B, C, F, and H), or a Welch's t-test (E). *, $p < 0.05$; **, $p < 0.01$; ***, $p < 0.001$; ns, not significant.

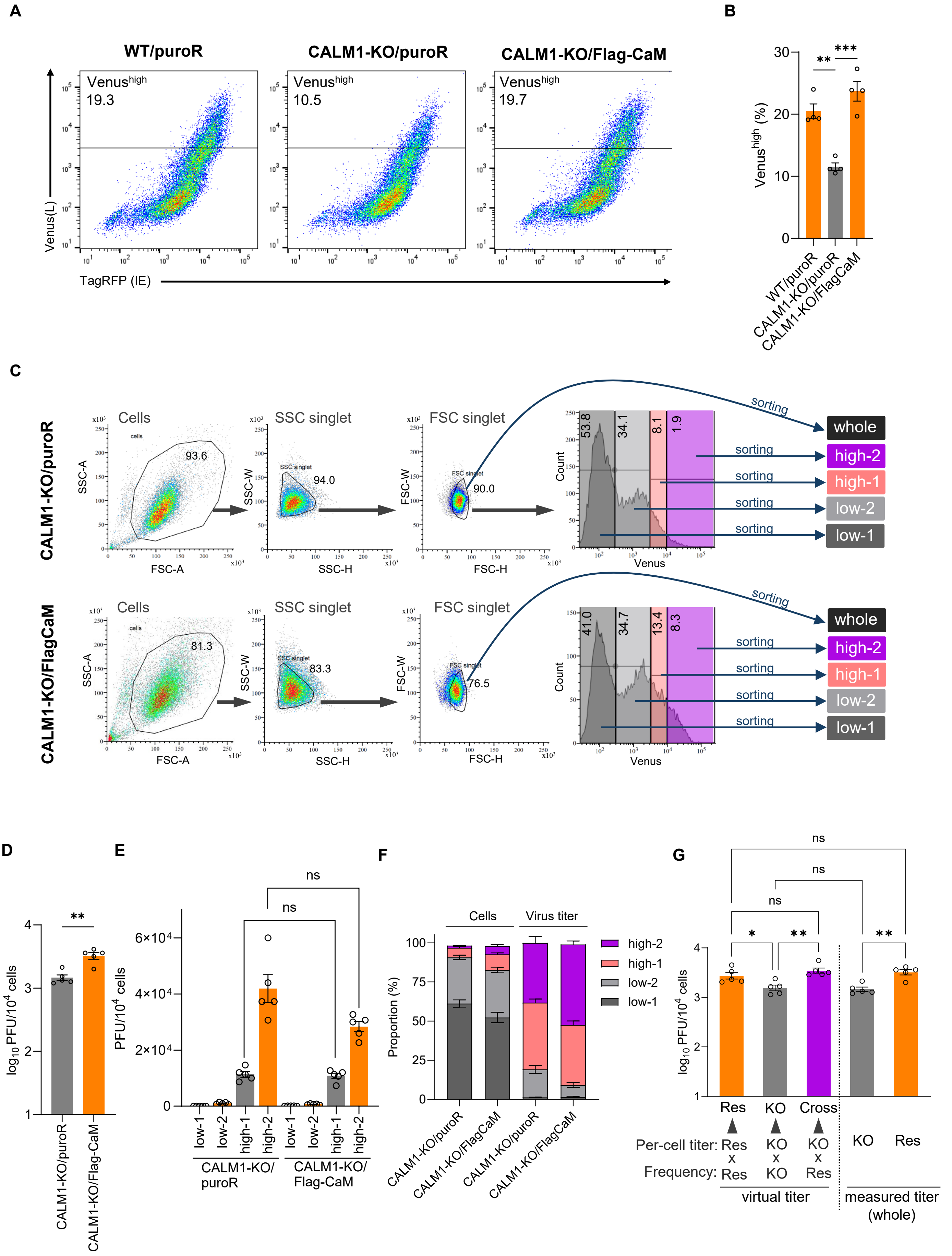


*(legend on next page)*

**S-Fig. 8 Y. Maruzuru et al.**

**S-Fig. 8. Complementation of CALM1 restores viral gene expression heterogeneity and progeny virus production.**

(A and B) Control (WT/puroR, CALM1-KO/puroR) and Flag-CaM complemented CALM1-KO (CALM1-KO/Flag-CaM) cells were infected with rICP47/vUs11 at an MOI of 5. At 12 h post-infection, cells were analyzed by flow cytometry. (A) Representative flow cytometry plots showing TagRFP (IE) and Venus (L) fluorescence. The gate for the Venus$^{high}$ population (virus-producing cells) is shown, with the percentage of cells within this gate indicated. (B) Quantification of the percentage of Venus$^{high}$ cells from (A).

(C to G) CALM1-KO/puroR (control) and CALM1-KO/Flag-CaM (rescue) cells were infected as in (A) and subjected to fluorescence-activated cell sorting (FACS) at 12 h post-infection.

(C) Gating strategy for cell sorting. Single cells (SSC- and FSC singlet) were sorted into the whole population or fractionated into four subpopulations (low-1, low-2, high-1, high-2) based on Venus (L) fluorescence intensity.

(D) Viral titers (PFU per $10^4$ cells) from the sorted "whole" singlet population.

(E) Viral titers of each of the four sorted subpopulations in (C) (i.e., the per-cell titers used in G).

(F) Stacked bar graphs showing, for CALM1-KO/Flag-CaM (Res) and CALM1-KO/puroR (KO) cells, the subpopulation composition ("Cells") and each subpopulation's contribution to the total viral yield ("Virus titer"), both expressed as percentages.

(G) Virtual titer analysis. For each Venus-defined subpopulation, the product of its frequency among infected cells ("Frequency") and its per-cell titer ("Per-cell titer") was calculated, and these products were summed across subpopulations to obtain the virtual titer. The source of each parameter (Res, CALM1-KO/Flag-CaM; or KO, CALM1-KO/puroR) is indicated below each bar: the "Res" and "KO" bars use the Frequency and Per-cell titer from the same cells, whereas the "Cross" bar combines the Res Frequency with the KO Per-cell titer. Measured whole-population titers (as in D) are shown on the right for comparison.

Each value represents the mean ± SE of four (B) or five (D to G) biological replicates. Statistical analyses were performed using one-way ANOVA followed by Tukey's multiple-comparison test (B, E, and G), or a Welch's t-test (D). *, $p < 0.05$; **, $p < 0.01$; ***, $p < 0.001$; ns, not significant.

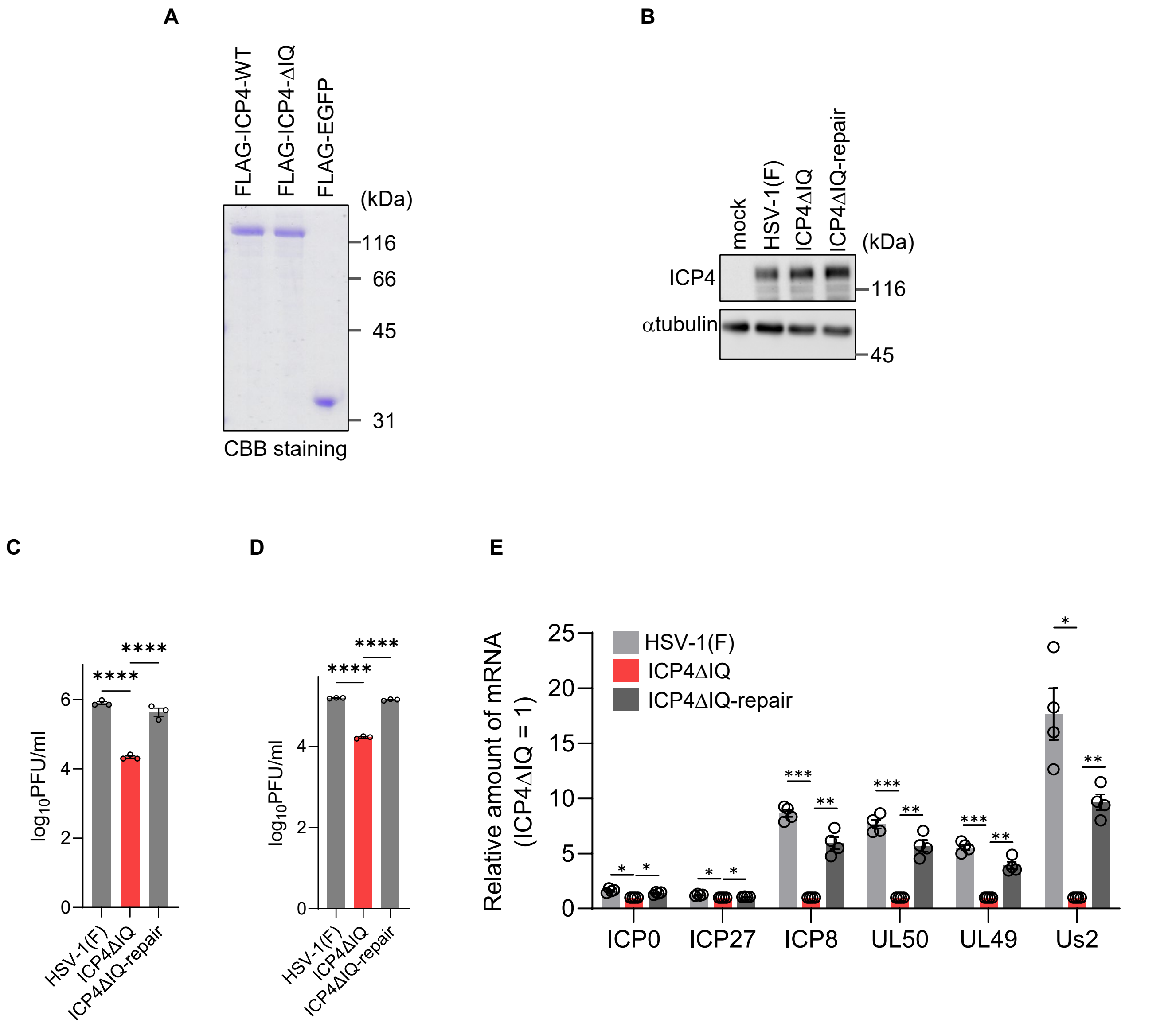


**S-Fig. 9. Deletion of the ICP4 IQ-like motif attenuates viral gene expression and replication.**
(A) Coomassie Brilliant Blue (CBB) staining of purified FLAG-tagged proteins (FLAG-ICP4-WT, FLAG-ICP4-ΔIQ, and FLAG-EGFP).
(B) HeLa cells were mock-infected or infected with wild-type HSV-1(F), ICP4ΔIQ, or ICP4ΔIQ-repair at an MOI of 5. Cell lysates were harvested at 6 h post-infection and analyzed by immunoblotting with antibodies against ICP4 and α-tubulin.
(C and D) HeLa cells were infected with wild-type HSV-1(F), ICP4ΔIQ, or ICP4ΔIQ-repair at an MOI of 5 for 12 h (C) or at an MOI of 0.05 for 48 h (D). Progeny virus yields were determined by plaque assay on Vero cells.
(E) HeLa cells were infected with wild-type HSV-1(F), ICP4ΔIQ, or ICP4ΔIQ-repair at an MOI of 5 for 6 h. Relative mRNA levels of IE (ICP0, ICP27), E (ICP8, UL50), and L (UL49, Us2) genes were quantified by qRT-PCR and normalized to the levels in ICP4ΔIQ-infected cells (set to 1).
Data in (C to E) are presented as mean ± SE of three (C and D) or four (E) biological replicates. Statistical analyses were performed using one-way ANOVA followed by Tukey's multiple-comparison test (C and D) or a one-sample t-test with Bonferroni correction (E). *, $p < 0.05$; **, $p < 0.01$; ***, $p < 0.001$; ****, $p < 0.0001$.

S-Fig. 9 Y. Maruzuru et al.

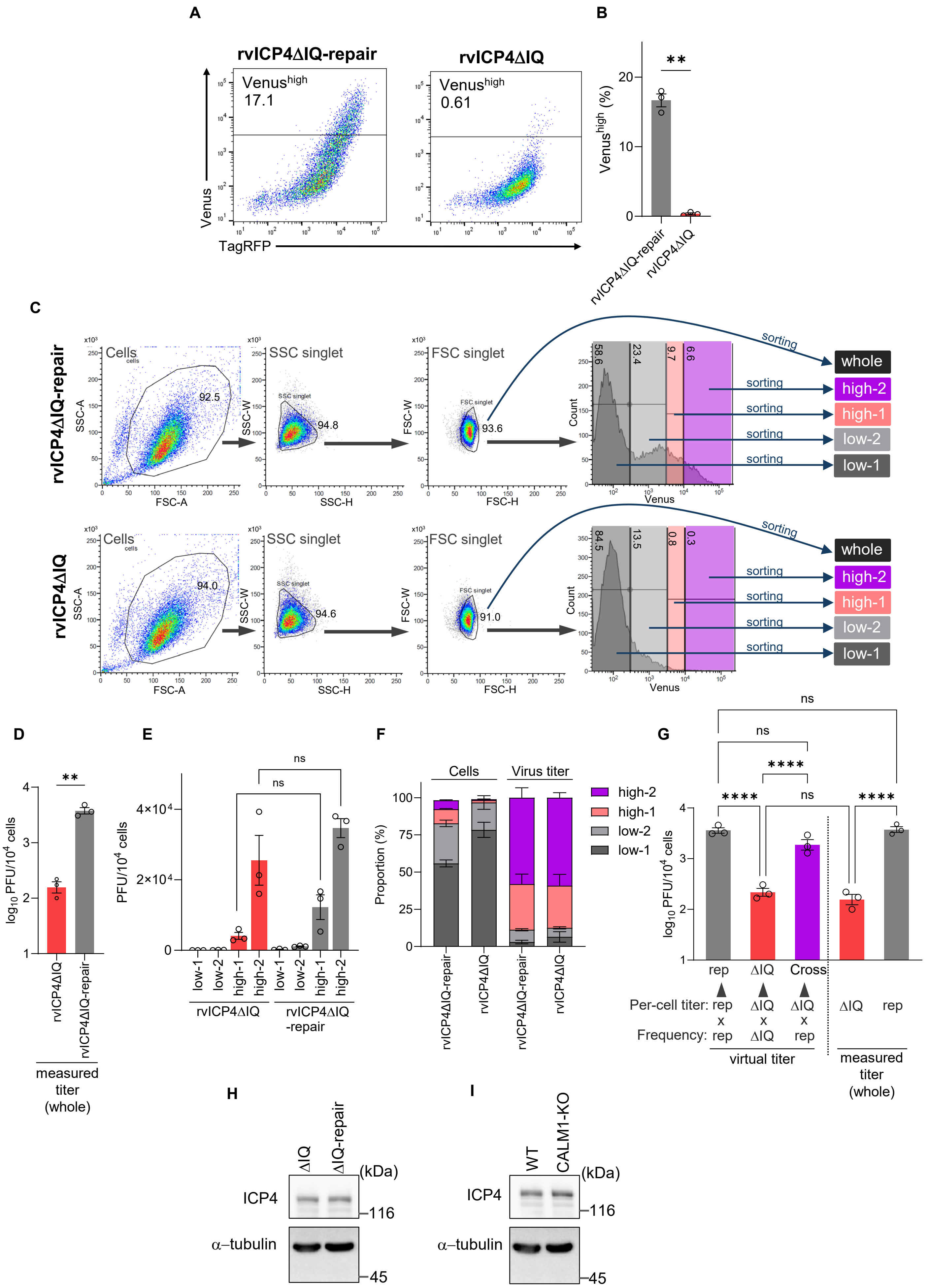


*(legend on next page )*

**S-Fig. 10 Y. Maruzuru et al.**

**S-Fig. 10. Deletion of the ICP4 IQ-like motif reduces progeny virus production by altering viral gene expression heterogeneity.**

(A and B) HeLa cells were infected with rICP47/vUs11/ICP4ΔIQ (rvICP4ΔIQ) or its repaired virus (rvICP4ΔIQ-repair) at an MOI of 5. At 12 h post-infection, cells were analyzed by flow cytometry. (A) Representative flow cytometry plots showing TagRFP (IE) and Venus (L) fluorescence. The gate for the Venus$^{high}$ population (virus-producing cells) is shown, with the percentage of cells within this gate indicated. (B) Quantification of the percentage of Venus$^{high}$ cells from (A).

(C to G) HeLa cells were infected as in (A) and subjected to fluorescence-activated cell sorting (FACS) at 12 h post-infection.

(C) Gating strategy for cell sorting. Single cells (SSC- and FSC singlet) were sorted into the whole population or fractionated into four subpopulations (low-1, low-2, high-1, high-2) based on Venus (L) fluorescence intensity for both rvICP4ΔIQ-repair and rvICP4ΔIQ infections.

(D) Viral titers (PFU per $10^4$ cells) from the sorted "whole" singlet population.

(E) Viral titers of each of the four sorted subpopulations in (C) (i.e., the per-cell titers used in G).

(F) Stacked bar graphs showing, for rvICP4ΔIQ-repair– and rvICP4ΔIQ-infected cells, the subpopulation composition ("Cells") and each subpopulation's contribution to the total viral yield ("Virus titer"), both expressed as percentages.

(G) Virtual titer analysis. For each Venus-defined subpopulation, the product of its frequency among infected cells ("Frequency") and its per-cell titer ("Per-cell titer") was calculated, and these products were summed across subpopulations to obtain the virtual titer. The source of each parameter (rep, rvICP4ΔIQ-repair; or ΔIQ, rvICP4ΔIQ) is indicated below each bar: the "rep" and "ΔIQ" bars use the Frequency and Per-cell titer from the same infection, whereas the "Cross" bar combines the rep Frequency with the ΔIQ Per-cell titer. Measured whole-population titers (as in D) are shown on the right for comparison.

(H) HeLa cells were infected with ICP4ΔIQ or ICP4ΔIQ-repair (MOI of 5). At 3.5 h post-infection, cell lysates were analyzed by immunoblotting with the indicated antibodies.

(I) WT or CALM1-KO cells were infected with wild-type HSV-1(F) (MOI of 5). At 3.5 h post-infection, cell lysates were analyzed by immunoblotting with the indicated antibodies.

Data are representative of three independent experiments (H and I). Each value represents the mean ± SE of three (B, D to G) biological replicates. Statistical analyses were performed using one-way ANOVA followed by Tukey's multiple-comparison test (E and G), or a Welch's t-test (B and D). *, $p < 0.05$; **, $p < 0.01$; ****, $p < 0.0001$; ns, not significant.
